# A framework for measuring national trends in the extinction risk of evolutionarily distinct species using the EDGE Index

**DOI:** 10.64898/2026.10.16.759618

**Authors:** Julie L.M. Campana, Sebastian Pipins, J. Alexander Nieto Lawrence, Steve Bachman, Ignacio Ramos-Gutiérrez, Hlengiwe Mtshali, Ditiro Moloto, Domitilla Claudia Raimondo, Claire Régnier, James Rosindell, Lize von Staden, Félix Forest, Roseli Pellens, Krystal A. Tolley, Rikki Gumbs, Guillaume Gigot

## Abstract

The biodiversity crisis is threatening evolutionarily distinct species, those that constitute long periods of unique evolutionary history or that are from highly imperilled lineages. The EDGE (Evolutionarily Distinct and Globally Endangered) index was designed to track the changing extinction risks of these species and is listed as an indicator in the Kunming-Montreal Global Biodiversity Framework. Here, we provide a set of guidelines for calculating the EDGE index at the national level and for addressing data uncertainty. We distinguish two approaches for calculating a country-level EDGE index, either generated directly by using national data or from disaggregated global data. Using two national Red Lists, we show the contrasting state of high evolutionarily distinct nesting birds across French territories, and the elevated threat status of high evolutionarily distinct endemic species for South African vertebrates. Using global extinction risk data, we then reveal the deteriorating state of birds, mammals, and amphibians over time, and we highlight the countries with the highest proportions of threatened evolutionarily distinct species. Application of these guidelines should aid countries in monitoring their evolutionarily irreplaceable species, which in turn can help target local conservation actions, while contributing to global efforts to preserve the tree of life.

## 1. Introduction

With the sixth mass extinction underway (Barnosky et al., 2011; Ceballos et al., 2015, 2017; Cowie et al., 2022), halting biodiversity loss has become an undeniable and urgent priority (Díaz, Settele, Brondízio, Ngo, Agard, et al., 2019). This crisis threatens all dimensions of biodiversity, from genes to ecosystems through species and the evolutionary lineages to which they belong (Mace et al., 2003; Díaz et al., 2020; Díaz, Settele, Brondízio, Ngo, Agard, et al., 2019). It is also leading to significantly negative impacts on Nature’s Contributions to People (NCP; Díaz et al., 2019) including deteriorating ecosystem services and quality of life (Díaz et al., 2020).

In 2022, the 15th Conference of Parties (COP) of the Convention on Biological Diversity adopted the Kunming-Montreal Global Biodiversity Framework (KM-GBF), which contains four overarching goals and 23 targets (GBF 2050 Goals; https://www.cbd.int/gbf/goals). Parties then adopted the Monitoring Framework (COP 16.2; CBD, 2025), defining a set of indicators to track progress towards these goals and targets in the coming decades. The effective application of these indicators at the national scale requires standardised methodologies and guidelines to aid countries in monitoring their national biodiversity in a way that can contribute towards global goals.

Following the recommendations of the Global Assessment on Biodiversity and Ecosystem Services (Díaz, Settele, Brondízio, Ngo, Guèze, et al., 2019), the KM-GBF adopted two tree of life indicators: the PD (Phylogenetic Diversity) indicator (Faith et al., 2018; Díaz, Settele, Brondízio, Ngo, Agard, et al., 2019) and the EDGE (Evolutionarily Distinct and Globally Endangered) Index (Gumbs, Chaudhary, et al., 2023). The rationale behind these indicators is that the conservation of the diversity within the tree of life is fundamental for preserving future options for humanity and Nature’s Contributions to People (Faith, 2016; Molina-Venegas, 2021; Forest et al., 2007; Díaz, Settele, Brondízio, Ngo, Guèze, et al., 2019), and for safeguarding the evolutionary breadth of all species (Faith, 1992a).

The tree of life is a metaphor used to describe how all extant and extinct species are connected through their evolutionary history. The more closely related species are, the more shared characters they will have. Phylogenetic diversity is a way to measure this diversity (Faith, 1992a), and it has been considered in conservation prioritisation studies for more than 30 years (Faith, 1992b; May, 1990; Vane-Wright et al., 1991; Pavoine et al., 2005; Forest et al., 2007; Pellens & Grandcolas, 2016). Species with greater amounts of unique evolutionary history will have more distinct combinations of characters and such species will contribute more to total phylogenetic diversity (Faith, 1992b, 1992a). Species that are the only, or the few, representatives of long branches of the tree of life are therefore irreplaceable because their extinction would result in the loss of distinct traits (*e.g.*, genes, morphology, behaviours) harboured only by these species. To safeguard the tree of life, the diversity of species and lineages that have emerged over evolutionary time must be conserved.

One of the strategies proposed to reduce the loss of evolutionarily unique species and lineages is the EDGE approach, which prioritises species that are both “Evolutionarily Distinct” (ED) and “Globally Endangered” (GE) (Isaac et al., 2007; Gumbs, Gray, et al., 2023). This approach has been applied to several taxonomic groups and has contributed to the conservation of more than 150 EDGE species across 50 countries by the Zoological Society of London’s EDGE of Existence program (https://www.edgeofexistence.org/). More recently, the approach has been adapted as a KM-GBF indicator, the “EDGE Index”, which measures the trends in the extinction risk of evolutionarily distinct species (CBD, 2022a, 2022b, 2025). In line with CBD reporting, parties are encouraged to report an EDGE Index at the national level. However, the approach adopted depends on the availability of data. When applied at the national level, the EDGE Index tracks changes in extinction risk status for threatened evolutionarily distinct species, supporting the protection of biodiversity nationally whilst also contributing to global efforts to preserve the tree of life (Gumbs, Chaudhary, et al., 2023).

Here, we propose a protocol for calculating the EDGE Index at the national level. We address two types of country-level EDGE Index compilations, the choice of which depends on the availability of national extinction risk data: (1) the National EDGE Index, which uses national extinction risk data, and (2) the Global EDGE Index, which uses global extinction risk data. Our protocol outlines the methodological process for (i) integrating national datasets, (ii) disaggregating data from global datasets if national data are not available, (iii) assessing uncertainty associated with these compilations, and (iv) estimating the data completeness thresholds required to obtain robust estimations. We then put these guidelines into practice by calculating the National EDGE Index for two biodiversity rich countries with different geographical configurations, France and South Africa, and exploring different spatial and temporal scales. For France, we focused on nesting birds in Metropolitan France and nine overseas territories. For South Africa, we assessed all jawed vertebrate groups and also examined trends over time in endemic and non-endemic lizards, snakes, tortoises, turtles and amphibians. We then calculated the Global EDGE Index for all jawed vertebrate groups across countries and we assessed the sensitivity of the country-level EDGE Index to data incompleteness.

## 2. EDGE Index disaggregated at a national level

### 2.1. The EDGE Index

The EDGE Index measures the conservation status of evolutionarily distinct species over time in a given spatial unit (country, state, ecoregion, etc.). This index, which builds on the EDGE2 protocol (Gumbs, Gray, et al., 2023), was developed by the IUCN SSC Phylogenetic Diversity Specialist Group (PDSG), and features in the KM-GBF as component indicator under the Goal A (CBD, 2025).

We focus on the first component of the EDGE Index (Gumbs, Gray, et al., 2023, Appendix S1) and reformulate it as “the proportion of threatened or extinct priority evolutionarily distinct species through time”. We define “priority evolutionarily distinct species” (hereafter, high ED species) as those with ED value above the median for a taxonomic group of interest, such as mammals or squamate reptiles (Gumbs, Chaudhary, et al., 2023). We define threatened species as those that are Critically Endangered (CR), Endangered (EN) or Vulnerable (VU) on the IUCN Red List; species considered extinct are those that are Critically Endangered Possibly Extinct (CR (PE)), Regionally Extinct (RE), Extinct in the Wild (EW), or Extinct (EX) on the IUCN Red List; species considered not threatened are those that are Least Concern (LC) or Near Threatened (NT) on the IUCN Red List. The EDGE Index is therefore calculated as a value ranging from 0, when no high ED species are threatened or extinct to 1, when all high ED species are either threatened with extinction or have gone extinct. The EDGE Index allows for the incorporation of extinction risk data at any spatial scale regarding the availability of Red List data, enabling countries (or other geographical units) to track the changing status of the high ED species found within their borders. In the following sections, we outline the procedure we recommend for calculating the country-level EDGE Index for a given taxonomic group, represented by the workflow of Figure 1.

**Figure 1.**
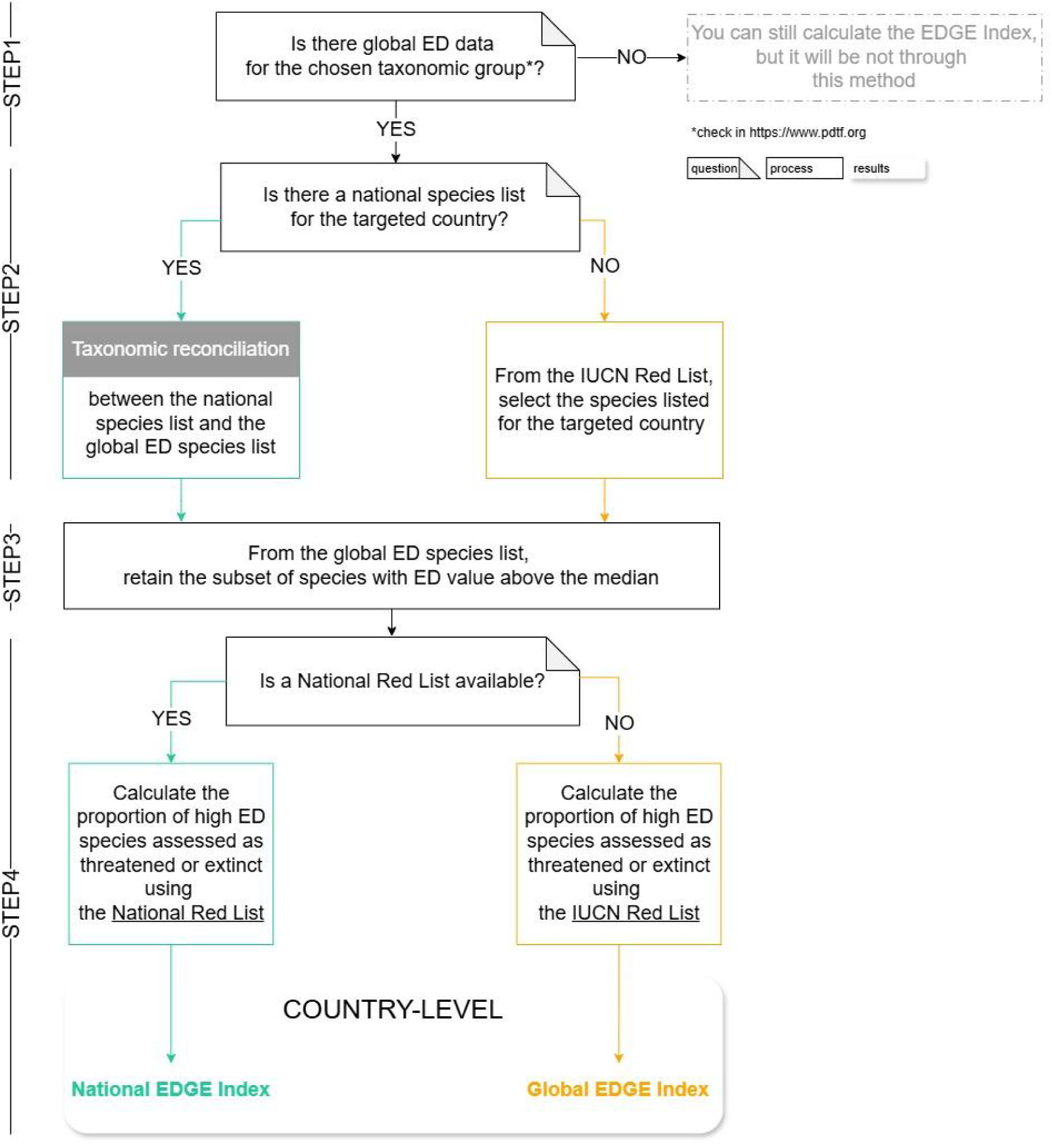
The recommended four-step procedure (and accompanying decisions) for calculating the country-level EDGE Index for a chosen taxonomic group for a targeted country or at any subglobal spatial scale based on the availability of Red List data. There are two versions of the country-level EDGE Index based on the availability of national extinction risk data: on the left, the blue path describes the compilation of the National EDGE Index, which combines global data of evolutionary distinctiveness and national extinction risk data (from the National Red List), and on the right, the orange path describes the compilation of the Global EDGE Index, which combines global data of evolutionary distinctiveness and global extinction risk (from the IUCN Red List).

### 2.2. The country-level EDGE Index calculation

#### 2.2.1. Data requirements

Two datasets are required to produce the country-level EDGE Index: (1) a global ED species list, with the set of species for a chosen taxonomic group with above median ED value; (2) a national species checklist for the chosen taxonomic group with native occurrences for a country or other spatial configuration/unit, as well as threat assessment data, usually based on the IUCN Red List.

The global ED species list is available from the PDSG’s EDGE Index platform (https://www.pdtf.org/), which compiled the lists of species for all major taxonomic groups that have been comprehensively assessed under the EDGE2 protocol (Gumbs, Gray, et al., 2023), which currently includes all jawed vertebrates (Gumbs et al., 2024), flowering plants (Forest et al., 2026), and crayfish (Pipins et al., 2026). The PDSG is committed to maintaining and updating global EDGE datasets for the next ten years.

#### 2.2.2. Guidelines for calculating the country-level EDGE Index

The four main steps, and accompanying decisions, for calculating the country-level EDGE Index are shown in Figure 1. This decision-making process depends on the availability of data, notably those relative to national extinction risk, given that the more comprehensive the national species list is, the more accurate the threat assessment will be.

<u>STEP 1</u> - Define the taxonomic group(s) of interest From the PDSG platform, select the taxonomic group(s) of interest to obtain the global ED species data for those sets of species. We encourage the selection of all taxonomic groups available in order to have the most complete view of threatened ED species to be nationally protected in priority.
<u>STEP 2</u> - Taxonomic reconciliation between national species list and global ED data As the species names can differ between different databases, it is necessary to compare and retain only one species name. Following IUCN standards for a comprehensively assessed group (IUCN, 2026), we recommend a congruence of at least 80% between the national species list and the global ED species list, using the standardised IUCN species identifiers where possible to make the match.
<u>STEP 3</u> - National high ED species list From the global ED species list, retain the subset of species with ED value above the median (Gumbs, Gray, et al., 2023). This results in the national high ED species list. We recommend determining the high ED species list separately for each taxonomic group as each will have a different median ED value.
<u>STEP 4</u> - Country-level EDGE Index For a given taxonomic group in a given country, the EDGE Index is the proportion of high ED species with data sufficient assessments (*i.e.*, not Data Deficient or unassessed - Not Evaluated) that are either threatened with extinction or that are extinct at a national level (or any sub-global spatial scale data of threat assessment) and at a given time (ones of the Red List data used).

For <u>STEP 2</u> and <u>3</u>, as mentioned above, a taxonomic match of 80% is desirable (IUCN, 2026), although a match as low as 50% will still produce reliable EDGE Index estimations as long as the taxonomic group in question has more than three high ED threatened species occurring within the country (see section 6.).

When available and comprehensive, we encourage the use of national extinction risk data to calculate the country-level EDGE Index (Figure 1, blue path). However, we do not suggest supplementing sub-global extinction risk data gaps with global ones because almost all Red List assessments are not aligned timing wise, the differing spatial scales of the two data types can make inferences unreliable and obscure national trends. When there is incomplete or unreliable or even no national extinction risk data available, we recommend that a country draws assessments exclusively from the IUCN Red list. Specifically for endemic species, national and global threat assessments are confounded for the majority of the taxonomic groups because of the unique spatial location of these species.

The outlined country-level EDGE Index calculation (<u>STEP 1</u> →<u>STEP 4</u>) can then be repeated for every time point for which there is a comprehensive set of extinction risk assessments.

In the absence of national extinction risk data, the country-level EDGE Index can be calculated by using IUCN Red List, which we have produced for all countries recognised by the IUCN (see section 6.), available <u>here</u>. Following terminology used for the Red List Index (RLI; Raimondo et al., 2023; Butchart et al., 2025) we refer to this version of the country-level EDGE Index as the Global EDGE Index (Figure 1, orange path), calculated as the proportion of assessed high ED species listed as threatened on the IUCN Red List for a given taxonomic group occurring in a specific territorial configuration.

#### 2.2.3. Calculating confidence intervals

Data Deficient (DD) and unassessed species (NE, Not Evaluated) are very common in most national Red Lists (Bland et al., 2017, 2015; Cazalis et al., 2023) and many taxonomic groups such as plants (Gallagher et al., 2023; Bachman et al., 2024). If not taken into consideration, the large number of DD/NE species could affect the reliability of a country-level EDGE Index. We suggest including confidence intervals around a country-level EDGE Index estimate that account for the uncertainty in the extinction risk of DD/NE species.

Following an approach used for the RLI (Raimondo et al., 2023), we recommend accounting for uncertainty by first assigning high ED species currently classified as DD/NE to threatened or non-threatened Red List categories using probabilities equal to the proportion of threatened high ED species (among assessed, data-sufficient species) within a given taxonomic group and for a given spatial level. This procedure should then be repeated 1,000 times, and uncertainty can be quantified using the 95% confidence interval across iterations.

### 2.3. Practical applications

The EDGE Index is used to inform how high ED species are faring across different countries and can be estimated over time if the group of species has been comprehensively assessed over multiple time points. We provide two examples from biodiversity rich countries: France and South Africa. We follow the four-step guidelines described above, corresponding to the blue path of Figure 1, to produce the National EDGE Index for different taxonomic groups. We then contrast it with the Global EDGE Index, using global extinction risk data corresponding to the same year, to highlight the differences between the two versions of the country-level EDGE Index and to show the added resolution of incorporating national data. We then produce the Global EDGE Index for jawed vertebrates across all countries using IUCN Red List data from 2023. We follow the four-step guidelines described above, corresponding to the orange path of Figure 1.

## 3. French EDGE Index for nesting birds across ten territories

French territories are distributed across different continents, oceans and in the Caribbean Sea, covering wide latitudinal and longitudinal ranges. They have important differences not only in species richness and threat status (Régnier & Gargominy, 2018; Table S1), but also in the amount of evolutionary history they represent. This fragmented territorial configuration provides an opportunity to explore the outcomes of the EDGE Index in different geographical and ecological contexts.

Nesting birds represent the taxonomic group with the highest coverage in both global and French threat assessments across a majority of territories (∼90% vs. ∼56%; Supplementary materials). Hence, ten French territories with a threat assessment were selected: three continental territories, Metropolitan France (including Corsica) in Europe, French Guiana in South America, and Adélie Land in Antarctic; and seven islands with Guadeloupe and Martinique in the Caribbean Sea, French Polynesia in the Pacific Ocean, Mayotte, Scattered Islands and La Réunion in the Indian Ocean, and Subantarctic Islands in the Austral Ocean (Figure S1). Respectively to the primary EDGE Index definition (Gumbs, Chaudhary, et al., 2023), we considered species identified as present and native from a given territory (Supplementary materials). For highly isolated islands (*e.g.*, Mayotte) where a subspecies was the only representative of the species in the site, we included the subspecies in the analysis. In such cases, the global ED value of the subspecies was considered equivalent to its parent (Table S2).

Following the procedure described above (Figure 1, left path), we calculated the two versions of the French EDGE Index (see section 2.3.) separately for each of the ten territories indicated above. For each version, we calculated the confidence intervals based on the number of DD/NE species relative to the Red List used for a given territory (see above).

The French National EDGE Index differs across the territories ranging from 0.18 for French Guiana to 0.60 for Scattered Islands and Adélie Land (Figure 2; Table 1). On average, the overseas territories have a higher National EDGE Index than Metropolitan France (0.45 *vs.* 0.34; Table 1). In the overseas territories, the National EDGE Index is similar within French Polynesia, Scattered Islands, and Adélie Land (0.58, 0.6 and 0.6) whereas these territories are located in different oceans (Pacific Ocean, Indian Ocean and Austral Ocean). Surprisingly, although adjacent in the Caribbean Sea, Guadeloupe and Martinique have very different National EDGE Index (0.49 and 0.38), as for Mayotte and La Réunion, which are relatively closed in the Indian Ocean, and have a different National EDGE Index (0.36 and 0.46).

**Figure 2.**
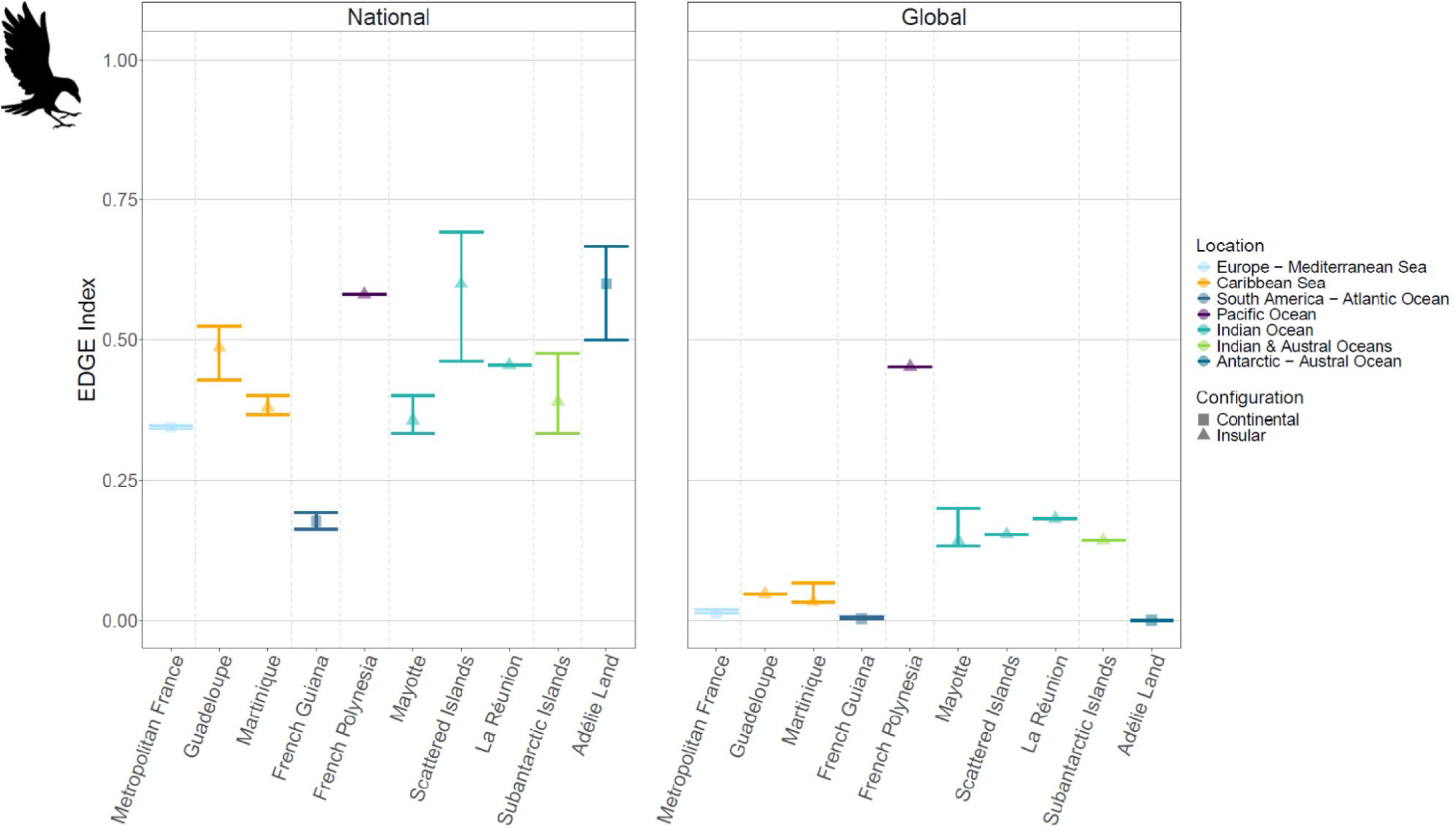
The French EDGE Index for nesting birds across the ten territories. Coloured dots represent the EDGE Index value and coloured bars represent the associated confidence intervals. Rectangles and triangles represent the EDGE Index value for territories with a continental and an insular configuration, respectively. “National” and “Global” panels represent respectively the values of the National EDGE Index (using National Red List) and the values of the Global EDGE Index (using IUCN Red List) for each territory.

**Table 1.** Summary of the French National EDGE Index results for the ten French territories calculated using local threat assessments. Configuration: indicates if the territory presents a continental *vs.* an insular topographic configuration; Year: date of the National Red List; N.ass sp: number of assessed species. N.sp DD/NE: number of species categorized as DD/NE; N.ED sp (% tot sp): number of high ED species (percentage relatively to the total number of species, *i.e.*, N.ass sp + N.sp DD/NE); N.ED sp and ass (% N.ass. sp): number of high ED species and assessed (percentage relatively to the number of assessed species, *i.e.*, N.ass sp); N.ED sp DD/NE (%N.sp DD/NE): number of high ED species categorized DD/NE (percentage relatively to the number of species categorized as DD/NE, *i.e.*, N.sp DD/NE); EDGE Index [confidence interval]: value of EDGE Index with its associated confidence interval calculated by using the number of DD/NE species relative to the National Red List.

| Territory | Configuration | Year | N.ass sp | N.sp DD/NE | N.ED sp<br>(% tot sp) | N.ED sp and ass<br>(% N.ass sp) | N.ED sp DD/NE<br>(% N.sp DD/NE) | EDGE Index<br>[confidence interval] |
| --- | --- | --- | --- | --- | --- | --- | --- | --- |
| Metropolitan France | Continental | 2016 | 276 | 2 | 152<br>(54.7%) | 151<br>(99.3%) | 1<br>(0.66%) | 0.344<br>[0.342, 0.349] |
| Guadeloupe | Insular | 2021 | 66 | 7 | 42<br>(57.5%) | 37<br>(88.1%) | 5<br>(11.6%) | 0.486<br>[0.429, 0.548] |
| Martinique | Insular | 2020 | 57 | 1 | 30<br>(51.7%) | 29<br>(96.7%) | 1<br>(3.33%) | 0.379<br>[0.367, 0.400] |
| French Guiana | Continental | 2017 | 494 | 61 | 312<br>(56.2%) | 276<br>(88.5%) | 36<br>(11.5%) | 0.178<br>[0.163, 0.192] |
| French Polynesia | Insular | 2015 | 56 | 0 | 31<br>(55.4%) | 31<br>(100%) | 0<br>(0%) | 0.581<br>[0.581, 0.581] |
| Mayotte | Insular | 2014 | 22 | 1 | 15<br>(65.2%) | 14<br>(93.3%) | 1<br>(6.67%) | 0.357<br>[0.333, 0.400] |
| Scattered Islands | Insular | 2015 | 12 | 8 | 13<br>(65%) | 10<br>(76.9%) | 3<br>(23.1%) | 0.600<br>[0.462, 0.692] |
| La Réunion | Insular | 2010 | 20 | 0 | 11<br>(55%) | 11<br>(100%) | 0<br>(0%) | 0.455<br>[0.455, 0.455] |
| Subantarctic Islands | Insular | 2015 | 36 | 9 | 21<br>(46.7%) | 18<br>(85.7%) | 3<br>(14.3%) | 0.389<br>[0.333, 0.476] |
| Adélie Land | Continental | 2015 | 7 | 1 | 6<br>(75%) | 5<br>(83.3%) | 1<br>(16.7%) | 0.600<br>[0.500, 0.667] |

The French National EDGE Index is on average 75.5% higher than the French Global EDGE Index, which reflects the greater proportion of high ED species threatened at a national scale (0.44 *vs.* 0.12 on average; Table S3). The variation is higher for the National EDGE Index, due to the higher proportion of ED species categorized as DD/NE at a national scale (8.81% *vs.* 1.16% on average; Table S3).

Irrespective of the Red List source, insular territories present on average higher French EDGE indices than continental territories (0.46 *vs.* 0.37 for National EDGE Index and 0.17 *vs.* 0.006 for Global EDGE Index; Table S3).

With its territories scattered across the globe, France presents a unique case study for the protection of phylogenetic diversity.

This study showed uneven vulnerability across the ten French territories, with the highest proportions of threatened high ED species in insular overseas territories. This “continents *vs.* islands” variability in the EDGE Index values can be explained by two main factors. First, island biodiversity was shown to be composed of many species pools representative of unique lineages across many branches of the tree of life (Véron et al., 2019; Warren et al., 2015, 2018). Beyond their evolutionary history characterized by spatial isolation (intimately linked to the geological history; Borregaard et al., 2017; Whittaker et al., 2008), the insular species are all the more vulnerable because of: (i) smaller population size, (ii) narrower ranges of distribution, and (iii) high rates of endemism (Myers et al., 2000; Kier et al., 2009; Tershy et al., 2015; Russell & Kueffer, 2019; Brook et al., 2008). Additionally, our results are consistent with the global trend of the intensification of bird extinction in islands over the last century (Sayol et al., 2021; Matthews et al., 2022), directly impacts phylogenetic diversity (Jetz et al., 2014) but also functional diversity (Matthews et al., 2024). These loss of diversity generally result from the loss of very specific and advantageous traits, such as the flight capacity, making insular birds unable to escape and therefore particularly vulnerable to various threats (Russell & Kueffer, 2019; Bellard et al., 2025), with invasive species (Bellard et al., 2025; Doherty et al., 2016) and climate change the two main emerging threats specifically (Russell & Kueffer, 2019 with data from Maxwell & et al., 2016 and Leclerc et al., 2018). Losing very unique bird species that contribute to very specific ecological functions (Sekercioglu et al., 2016; Pigot et al., 2020) can lead to significant disturbances (Anderson et al., 2011) in an insular ecosystem already highly fragile and sensitive to any environmental changes. However, these relevant studies remain in a global framework and do not allow the identification of the specific threats occurring in very small territories such as islands, their specific typology (archipelago *vs.* unique island, atoll *vs.* volcanic, etc.) potentially adding another dimension. Therefore, it is all more important to locally assess the threat levels given that a country can present extremely diverse environmental contexts as those that make up France’s territorial configuration and at any sub-spatial level such as localities in Metropolitan France.

Overall, France provides an interesting practical application across space of the disaggregation of the EDGE Index at a national level. Here, we focused on nesting species of birds since “nesting” is the consensual unit used to define standardised criteria across countries for biodiversity conservation, and therefore to be the most relevant for calculating the EDGE Index for a given territory. However, a significant proportion of bird species cross national administrative boundaries at different times of the year, often according to seasonal migration phases. Considering a pool of species according to their presence during the nesting season only tends to underestimate the proportion of high ED species locally threatened because the species present for a considerably short period (sometimes only a few hours for migratory species) on a given territory are ignored. For example, the vagrant species are all the more vulnerable if they are in their breeding season or if the territory represents an essential strategic stopover in their migration journey. The Global EDGE Index enables monitoring across the full distributions of the species but underestimates significantly the real proportion of high ED species threatened at a national level (see above). It would be interesting to investigate further the environmental ubiquity of these species directly linked to their ecological niche characteristics to establish an “(environmental) vulnerability criterion”. Indeed, if these specialist species are threatened and just passing through a given territory with specific environments, they could be highly vulnerable. This dilemma is quite tricky to resolve and highlights the importance of national assessments as a complement to global assessments to find the best methodological compromise leading to the more realistic estimation of the threat proportion at a local level. Ideally, and where possible, any species present in a territory, even for an extremely short period of time, should benefit from protective measures in proportion to the length of time the species is present in the territory. This more inclusive and conservative approach would be a good way to pool efforts in biodiversity preservation (Obura et al., 2021) and, in particular, to define measures to protect species within national borders while contributing to global efforts to preserve the tree of life as a whole.

Across the ten territories, the French National EDGE Index values were calculated over a time period from 2010 (La Réunion; Table 1) to 2021 (Guadeloupe; Table 1). Strengthening efforts to assess local threats would allow for precise monitoring of the effectiveness of protection measures implemented specifically in each French territory. The fragmented geographical distribution across the globe presents a real challenge. However, the coordinated teamwork between public institutions, associations, and scientific research allows progress to be made, as is the case for other taxonomic groups (*e.g.*, reptiles of Metropolitan France or vascular plants of La Réunion with two assessments), and represents a real lever for actions in the protection of French biodiversity.

## 4. South African EDGE Index for jawed vertebrates, with a particular attention to endemic amphibians, lizards & snakes, tortoises & turtles

Following the procedure described above (Figure 1, left path), we used national and global Red List assessments to calculate the two versions of the country-level EDGE Index (see section 2.3.) and associated confidence intervals for amphibians (assessed in 2025), birds (2015), lizards & snakes (lepidosaurs; 2022), mammals (2016), ray-finned fish (Actinopterygii; 2017), and tortoises & turtles (Testudines; 2022) for all species and for endemics only. Assessments for endemic species are equivalent in the national and global Red List data but differ for non-endemics. For comparison across taxonomic groups, we calculate unique values of the EDGE Index with the most recent National and global IUCN Red List assessments for each taxonomic group.

We also assessed trends in the National EDGE Index for the groups of lizards & snakes and tortoises & turtles between 1990 and 2022, and amphibians between 2004 and 2025. The trends were assessed for endemic and non-endemics species combined, and also for endemics only. For species endemic to South Africa, global and national assessments are functionally equivalent, as both are produced largely by the same South African assessors and institutes. Extinction risk data for these species were therefore obtained from the IUCN Red List.

Across all high ED South African vertebrates, the range of the National EDGE Index varies from 0.03 in lizards & snakes to 0.40 in ray-finned fish (Figure 3A; Table 2). For most vertebrate groups, the National EDGE Index is generally higher than the Global EDGE Index (Figure 3A). In some cases, the differences are marginal (*e.g.*, lizards & snakes, birds, mammals) while others show marked differences between approaches (*e.g.*, ray-finned fish - National EDGE Index = 0.40, Global EDGE Index = 0.278). In contrast, for tortoises & turtles the South African Global EDGE Index was higher than the National EDGE Index, reflecting the national assignment of LC to many species that do not have published global assessments (*Kinixys spekii*, *Kinixys zombensis*, *Pelomedusa subrufa*, *Pelusios sinuatus*, *Psammobates oculifer*), making the proportion of threatened species higher. Furthermore, the confidence intervals are smaller for some groups for South African National EDGE indices due to the lower number of DD/NE species. For example, 2% nationally *vs.* 13% globally for amphibians, and 4% nationally *vs.* 22% globally for ray-finned fish. However, this is not always the case, as a large proportion of high ED bird species have not been evaluated at a national level (9%).

**Figure 3.**
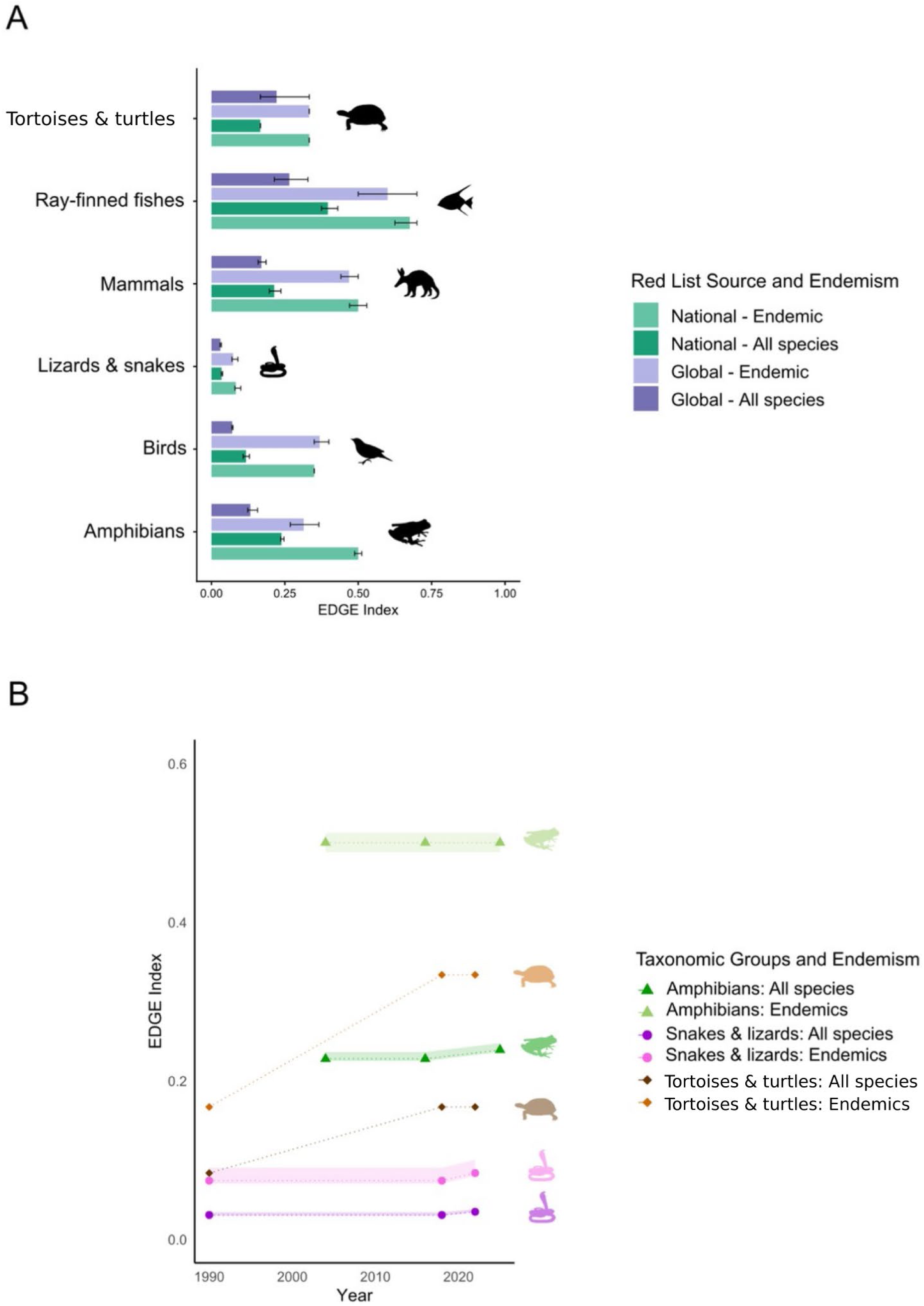
A: EDGE indices estimated for vertebrate groups from South Africa that have been comprehensively assessed for the IUCN Red List. EDGE indices were estimated based on the set of available globally disaggregated assessments (purple) and for the set of nationally produced assessments (green) for both endemics and for all species (endemics plus non-endemics). Error bars account for Data Deficient (DD) and Not Evaluated (NE) species by randomly resampling their threat category across 1,000 iterations. The figure indicates single time points of assessments, the years of which for each taxonomic group are: 2015 for birds, 2016 for mammals, 2017 for ray-finned fishes (Actinopterygii), 2022 for tortoises & turtles (Testudines) and lizards & snakes (lepidosaurs), and 2025 for amphibians. B: EDGE indices estimated for amphibians, lizards & snakes (lepidosaurs), and tortoises & turtles (Testudines) over time, based on multiple IUCN Red List assessments undertaken at the national level. The time periods are: 2004-2016-2025 for amphibians, 1990–2018-2022 for lizards & snakes (lepidosaurs) and tortoises & turtles (Testudines). EDGE indices at each assessment period are shown, joined by dotted lines to visualise the level of change over time. Points showing “all species” includes endemics plus non-endemics. Error margins are indicated as the solid ribbons using the same iterative approach with DD/NE species.

**Table 2.** A) Summary of South African National EDGE Index results for jawed vertebrates, stratified by endemism and the source of Red List threat categories: national *vs.* global categories. B) Summary of South African National EDGE Index results for herpetofauna across different assessment periods, specifically, amphibians, lizards & snakes (lepidosaurs), and tortoises & turtles (Testudines). Year: date of the National Red List; N.ass sp: number of assessed species. N.sp DD/NE: number of species categorized as DD/NE; N.ED sp (% tot sp): number of high ED species (percentage relatively to the total number of species, *i.e.*, N.ass sp + N.sp DD/NE); N.ED sp and ass (% N.ass. sp): number of high ED species and assessed (percentage relatively to the number of assessed species, *i.e.*, N.ass sp); N.ED sp DD/NE (%N.sp DD/NE): number of high ED species categorized DD/NE (percentage relatively to the number of species categorized as DD/NE, *i.e.*, N.sp DD/NE); EDGE Index [confidence interval]: value of EDGE Index with its associated confidence interval calculated by using the number of DD/NE species relative to the National Red List.

| Taxonomic group | Year | N.ass sp |  | N.sp DD/NE |  | N.ED sp (% tot sp) |  | N.ED sp and ass (% N.ass sp) |  | N.ED sp DD/NE (% N.sp DD/NE) |  | EDGE Index [confidence interval] |  |
| --- | --- | --- | --- | --- | --- | --- | --- | --- | --- | --- | --- | --- | --- |
|  |  | All species | Endemic | All species | Endemic | All species | Endemic | All species | Endemic | All species | Endemic | All species | Endemic |
| Amphibians | 2025 | 133 | 67 | 2 | 2 | 89<br>(65.9%) | 41<br>(59.4%) | 88<br>(98.9%) | 40<br>(97.6%) | 1<br>(1.1%) | 1<br>(2.4%) | 0.239<br>[0.236, 0.247] | 0.5<br>[0.488, 0.512] |
| Lizards-Snakes | 2022 | 378 | 142 | 7 | 6 | 237<br>(61.6%) | 100<br>(67.6%) | 232<br>(97.9%) | 96<br>(96%) | 5<br>(2.1%) | 4<br>(4%) | 0.034<br>[0.034, 0.038] | 0.083<br>[0.08, 0.1] |
| Mammals | 2016 | 314 | 51 | 31 | 6 | 228<br>(66.1%) | 34<br>(59.6%) | 201<br>(88.2%) | 32<br>(94.1%) | 27<br>(11.8%) | 2<br>(5.9%) | 0.214<br>[0.197, 0.237] | 0.5<br>[0.471, 0.529] |
| Tortoises-Turtles | 2022 | 19 | 7 | 0 | 0 | 12<br>(63.2%) | 6<br>(85.7%) | 12<br>(100%) | 6<br>(100%) | 0<br>(0%) | 0<br>(0%) | 0.167<br>[0.167, 0.167] | 0.333<br>[0.333, 0.333] |
| Ray-finned fish | 2017 | 113 | 56 | 5 | 3 | 72<br>(61%) | 40<br>(67.8%) | 68<br>(94.4%) | 37<br>(92.5%) | 4<br>(5.6%) | 3<br>(7.5%) | 0.397<br>[0.375, 0.431] | 0.676<br>[0.625, 0.7] |
| Birds | 2015 | 732 | 38 | 114 | 0 | 518<br>(61.2%) | 20<br>(52.6%) | 441<br>(85.1%) | 20<br>(100%) | 77<br>(14.9%) | 0<br>(0%) | 0.118<br>[0.108, 0.129] | 0.35<br>[0.35, 0.35] |

| Taxonomic group | Year | N.ass sp |  | N.sp DD/NE |  | N.ED sp<br>(% tot sp) |  | N.ED sp and ass<br>(% N.ass sp) |  | N.ED sp DD/NE<br>(% N.sp DD/NE) |  | EDGE Index<br>[confidence interval] |  |
| --- | --- | --- | --- | --- | --- | --- | --- | --- | --- | --- | --- | --- | --- |
|  |  | All species | Endemic | All species | Endemic | All species | Endemic | All species | Endemic | All species | Endemic | All species | Endemic |
| Amphibians | 2004 | 133 | 67 | 2 | 2 | 89<br>(65.9%) | 41<br>(59.4%) | 88<br>(98.9%) | 40<br>(97.6%) | 1<br>(1.1%) | 1<br>(2.4%) | 0.239<br>[0.236, 0.247] | 0.5<br>[0.488, 0.512] |
|  | 2016 | 133 | 67 | 2 | 2 | 89<br>(65.9%) | 41<br>(59.4%) | 88<br>(98.9%) | 40<br>(97.6%) | 1<br>(1.1%) | 1<br>(2.4%) | 0.239<br>[0.236, 0.247] | 0.5<br>[0.488, 0.512] |
|  | 2025 | 133 | 67 | 2 | 2 | 89<br>(65.9%) | 41<br>(59.4%) | 88<br>(98.9%) | 40<br>(97.6%) | 1<br>(1.1%) | 1<br>(2.4%) | 0.239<br>[0.236, 0.247] | 0.5<br>[0.488, 0.512] |
| Lizards<br>-Snakes | 1990 | 378 | 142 | 7 | 6 | 237<br>(61.6%) | 100<br>(67.6%) | 230<br>(97%) | 95<br>(95%) | 7<br>(3%) | 5<br>(5%) | 0.03<br>[0.034, 0.034] | 0.074<br>[0.07, 0.09] |
|  | 2018 | 378 | 142 | 7 | 6 | 237<br>(61.6%) | 100<br>(67.6%) | 230<br>(97%) | 95<br>(95%) | 7<br>(3%) | 5<br>(5%) | 0.03<br>[0.03, 0.034] | 0.074<br>[0.07, 0.09] |
|  | 2022 | 378 | 142 | 7 | 6 | 237<br>(61.6%) | 100<br>(67.6%) | 232<br>(97.9%) | 96<br>(96%) | 5<br>(2.1%) | 4<br>(4%) | 0.034<br>[0.034, 0.038] | 0.083<br>[0.08, 0.1] |
| Tortoises<br>-Turtles | 1990 | 19 | 7 | 0 | 0 | 12<br>(63.2%) | 6<br>(85.7%) | 12<br>(100%) | 6<br>(100%) | 0<br>(0%) | 0<br>(0%) | 0.083<br>[0.083, 0.083] | 0.167<br>[0.167, 0.167] |
|  | 2018 | 19 | 7 | 0 | 0 | 12<br>(63.2%) | 6<br>(85.7%) | 12<br>(100%) | 6<br>(100%) | 0<br>(0%) | 0<br>(0%) | 0.167<br>[0.167, 0.167] | 0.333<br>[0.333, 0.333] |
|  | 2022 | 19 | 7 | 0 | 0 | 12<br>(63.2%) | 6<br>(85.7%) | 12<br>(100%) | 6<br>(100%) | 0<br>(0%) | 0<br>(0%) | 0.167<br>[0.167, 0.167] | 0.333<br>[0.333, 0.333] |

The South African National EDGE Index tracked over time shows a gradual worsening trend for both amphibians, lizards & snakes, and tortoises & turtles since 1990 (Figure 3B). For all taxonomic groups, the endemic species have notably higher EDGE indices than for all species (endemics plus non-endemics). Among lizards & snakes and amphibians, the National EDGE indices have not notably changed over time (0 for endemic amphibians and 0.01 for all amphibians, all lizards & snakes, and endemic lizards & snakes). The slightly higher changes in National EDGE Index for all amphibians (endemics plus non-endemics) can be attributed to the genuine change in threat status for *Amietia hymenopus*. The National EDGE Index for tortoises & turtles shows a substantial increase between 1990 and 2018, where the National EDGE Index for all species doubles from 0.083 to 0.167. Endemic tortoises & turtles show a similar increase, from 0.167 to 0.333. This stark change is due to six species, previously classified as LC or NT, that experienced a shift to the threatened categories of VU or EN. Overall, for lizards & snakes and amphibians, it appears that the drivers of extinction risk were already well-established decades ago, but for tortoises & turtles, there has been a shift toward threatened categories since 2018.

South Africa is one of the world’s mega diverse countries, with endemism reaching 50% for reptiles, amphibians and freshwater fishes, and nearly 70% for plants (Skowno et al., 2019; Raimondo et al., 2023). These taxonomic groups, along with the birds, mammals and certain invertebrates have been comprehensively assessed for their IUCN Red List threat status both nationally and globally (Skowno et al., 2019; SANBI, 2025). Lizards & snakes, tortoises & turtles, and amphibians have comprehensively been assessed multiple times (Measey et al., 2019; Tolley et al., 2019, 2023) allowing not only status but also trends of biodiversity losses, or gains, to be tracked. The comprehensive sets of assessments, together with available global phylogenetic trees for vertebrates (Gumbs et al., 2024) places South Africa in a unique position to report on the EDGE Index to meet global reporting commitments on nearly all native vertebrates, and also to appraise its vast complement of endemic species.

The compilation of EDGE Index presented here following the guidelines outlined above for a suite of endemic South African vertebrate species that have been comprehensively assessed at least once shows that the National EDGE Index is higher (based on national assessments) than the Global EDGE Index (based on global assessments) (Figure 3A). Furthermore, the confidence intervals are generally smaller for the national datasets. This disparity is due to a large number of species that are categorised as DD/NE in the IUCN Red List but have been assessed nationally. Comparatively, the higher EDGE indices and the lower confidence intervals suggest that the national datasets are more sensitive to threat status and are more reliable in producing the most accurate EDGE Index for a taxonomic group.

Three major vertebrate groups - lizards & snakes, tortoises & turtles, and amphibians, all with approximately 50% endemism - have been assessed at multiple time points (Minter et al., 2004; Measey, 2011; Measey et al., 2019; Tolley et al., 2019, 2023) allowing the EDGE Index to be estimated over time for these groups (Figure 3B). Endemic EDGE amphibians are the most threatened group, with their EDGE Index indicating that half of all evolutionarily distinct endemic amphibians are threatened with extinction. The EDGE Index across all vertebrate groups shows that endemics are proportionally faring worse than all native species combined. However, for lizards & snakes and amphibians, the EDGE Index has worsened only marginally. Thus, the earliest assessments already show that high ED species were at risk, but few additional non-threatened high ED species have moved into threatened categories since that time. This suggests that the drivers of extinction risk were already well-established decades ago, but the severity of these threats may still have increased during this period, *i.e.*, uplistings between threat categories. In contrast, for tortoises & turtles, the EDGE Index has shown an increase over time, and this increase would ostensibly be due to both endemics and non-endemic species becoming more threatened over time. Although the increase is concerning, the change in the EDGE Index is 0.084 for all native species and 0.166 for endemics.

Given that the EDGE Index has either remained fairly stable over time, if not increasing in the case of tortoises & turtles, this suggests a lack of amelioration in threat status for South Africa’s most evolutionarily diverse reptiles and amphibians (Table S4). The lack of improvement of the EDGE Index demonstrates the critical need to advance targeted conservation efforts for species most at risk and evolutionarily distinct.

## 5. Global EDGE Index for jawed vertebrates

To set a benchmark for how many high ED species are threatened across countries at a global level, we calculated disaggregated global EDGE indices for all jawed vertebrates using spatial information from the IUCN Red List (2023.2). The Global EDGE Index was calculated using native ranges for all jawed vertebrate species and for each taxonomic group individually in each of the 251 spatial units with IUCN country-codes (hereafter, country). To do this, we combined a global list of high ED species with categories from the IUCN Red List (2023.2) for the following vertebrate taxonomic groups: amphibians, birds, mammals, lizards & snakes (lepidosaurs), tortoises & turtles (Testudines), crocodylians, sharks & rays (Chondrichthyes), and ray-finned fish (Actinopterygii). For each country and taxonomic group, both the EDGE Index and the number of high ED species (‘N’) were reported.

We found that island countries recurrently emerge as top-ranked hotspots of threatened high ED species. When considering the Global EDGE Index across all jawed vertebrates, Madagascar ranked highest (Figure S2), with 0.235 of its 2,046 high ED species being threatened. For taxonomic group-specific EDGE indices (Figure S3), the countries with the highest scores include French Polynesia for birds (0.389, N = 54), Jamaica for amphibians (0.900, N = 10), New Caledonia for lizards & snakes (0.737, N = 58), and Madagascar for mammals (0.518, N = 146). In 25 countries, all high ED crocodylians are threatened, each with one or two species, and all high ED tortoises & turtles are threatened in 134 countries, with the most speciose being Vietnam (N = 20). Tortoises & turtles have the highest mean EDGE Index across countries (0.892), followed by crocodylians (0.719) and sharks & rays (0.717).

For taxonomic groups with multiple global assessments, it is also possible to reveal how the status of high ED species have changed over time. We therefore assessed trends in the EDGE Index using information from the RLI, which tracks genuine changes in the Red List categories of species (Butchart et al. 2007). We assessed how the threat status of high ED species changed from first assessment to the most recent assessment available for the three taxonomic groups of jawed vertebrates with available RLI data (amphibians, birds, and mammals), reporting the changes in EDGE Index in each country.

When examining the mean change in the Global EDGE Index across countries (Figure 4), we found a slight average decrease for mammals (−0.060), and slight average increases for amphibians and birds (+0.019 and +0.008, respectively). In mammals, the improvement was mainly driven by countries with few high ED species harbouring the Humpback Whale, *Megaptera novaeangliae*, which was downlisted to non-threatened status between 1996 and 2008. Amphibians were the taxonomic group that showed the largest change in the number of threatened high ED species. For instance, Malaysia showed the largest decrease in the Global EDGE Index for amphibians (−0.071), due to the downlisting of ten high ED threatened endemic species between 1980 and 2022. However, Malaysia was ranked fourth highest for the increase in its Global EDGE Index for mammals (+0.033) and the seventh highest for its increased Global EDGE Index for birds (+0.027). Elsewhere, there were notable deteriorations, such as in Italy (+0.261), due to the uplisting of four endemic and one non-endemic species to a threatened status, and Chile (+0.182), where the Global EDGE Index increased to 0.591 due the uplisting of four high ED species.

**Figure 4.**
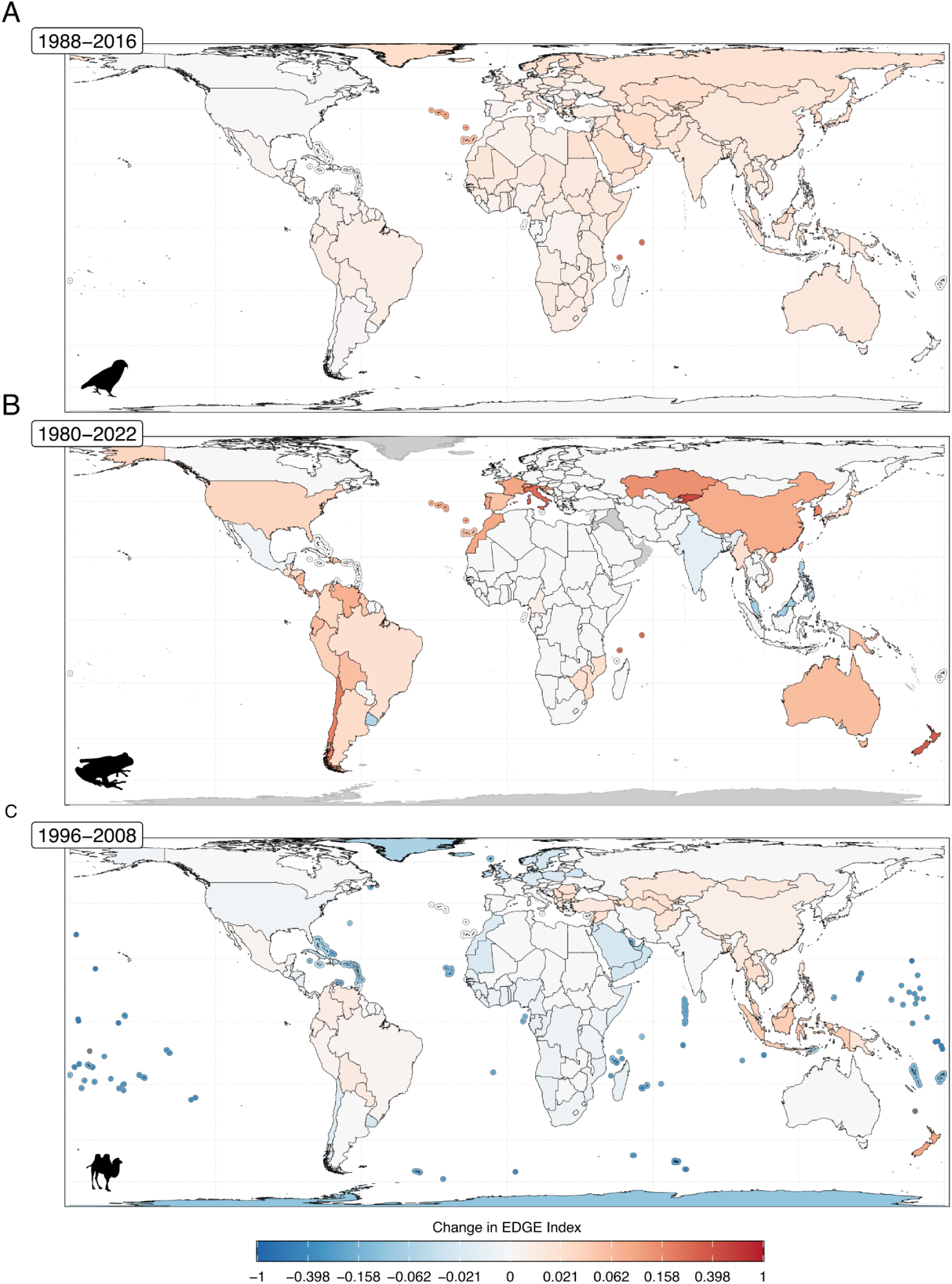
Trends in the Global EDGE Index for A: birds, B: amphibians, and C: mammals. Timepoints in the top left corner indicate the first and most recent years of global extinction risk assessment for each taxonomic group. Values vary between 1, where all high ED species moved from a non-threatened to a threatened IUCN Red List category, and −1 where all high ED species moved from a threatened category to a non-threatened category during the specified time period. Areas in grey in panel B indicate regions with no high ED amphibian species present.

Several of the regions highlighted here for their high Global EDGE indices align with known regions of highly concentrated threatened evolutionary history (Pipins et al., 2024), such as the Pacific islands for birds (Cooke et al., 2023), the Caribbean for amphibians (Luedtke et al., 2023), and the Mediterranean for sharks (Ferretti et al., 2013). Any minor reduction in the Global EDGE Index in one of these high scoring regions would therefore signify a noteworthy improvement in the status of the tree of life. However, there are also countries that stand out that are not usually distinguished for their high phylogenetic diversity. Italy, for example, was highlighted as a country of particular concern for amphibians, having experienced one of the largest increases in the amphibian Global EDGE Index, due to the uplisting of five high ED species to threatened status. This was largely due to the deteriorating state of its endemic and phylogenetically distinctive cave salamanders (Speleomantes *sp.*), many of which are predicted to undergo further climate-induced range contractions (Coppari et al., 2024; Iannella et al., 2025). Despite having the largest concentration of threatened amphibians in Europe, Italy has no Key Biodiversity Areas that are triggered by amphibians and contains none of the priority landscapes identified by the Global Amphibian Assessment (Re:wild, Synchronicity Earth, IUCN SSC Amphibian Specialist Group, 2023). By considering the threatened status of Italy’s high ED amphibians, the Global EDGE Index helps to highlight Italy as a global cause for concern. This demonstrates the utility of the Global EDGE Index, which is informative in revealing patterns across countries and for particular taxonomic groups, although it lacks the local context and national policy relevance of the National EDGE Index.

## 6. EDGE Index robustness

As not every species has a national or global extinction risk assessment, we sought to determine how robust the EDGE Index is to different degrees of data incompleteness. To do this, we used a rarefaction procedure on our global bird dataset, which involved incrementally increasing the proportion of species withheld from the EDGE Index calculation. We performed this analysis using Global EDGE indices in all countries with more than ten bird species. Using this approach, national species lists were rarified at random in 0.05 proportional increments (*i.e.*, rarefaction levels), up to 0.75, with each step repeated 1,000 times to incorporate uncertainty. At each rarefaction level, the EDGE Index was recalculated, and error was measured as percentage change from the EDGE Index of the complete data set. At each rarefaction level in each country, median and mean EDGE Index error values were computed from the 1,000 repeats. The relationship between rarefaction level and EDGE Index error was modelled using a generalised additive mixed model with a gamma distribution and log-link function.

We found that rarefaction level had a strong, positive, and linear effect on EDGE Index error (β = 1.21, t = 20.57, p < 0.001; Table S5). Mean EDGE Index error remained within ±6% up until a rarefaction level of 0.5 (Figure S4). This suggests that EDGE Index is reliably able to capture patterns in the extinction risk of high ED species in datasets with as low as 50% data coverage, making the EDGE Index a fit-for-purpose indicator for the changing status of the tree of life.

In addition, we found that the number of high ED threatened species in a country was a significant predictor of the EDGE Index error (F = 218.9, estimated degrees of freedom = 6.784, p < 0.0001; Figure 5). We observed a general stabilisation of this index in countries with more than three high ED threatened species. A mean of 93% of these countries maintained median EDGE Index errors within ±10% through rarefaction levels up to 0.5. This was over three times higher than in countries with three or fewer high ED threatened species (28%). This finding can be explained by the highly asymptotic relationship between the EDGE Index error and the number of high ED threatened species; if a nation contains one hundred species, and only one is a high ED threatened species, then even small changes in the extent of data completion would have a high chance of producing an inflated EDGE Index.

**Figure 5.**
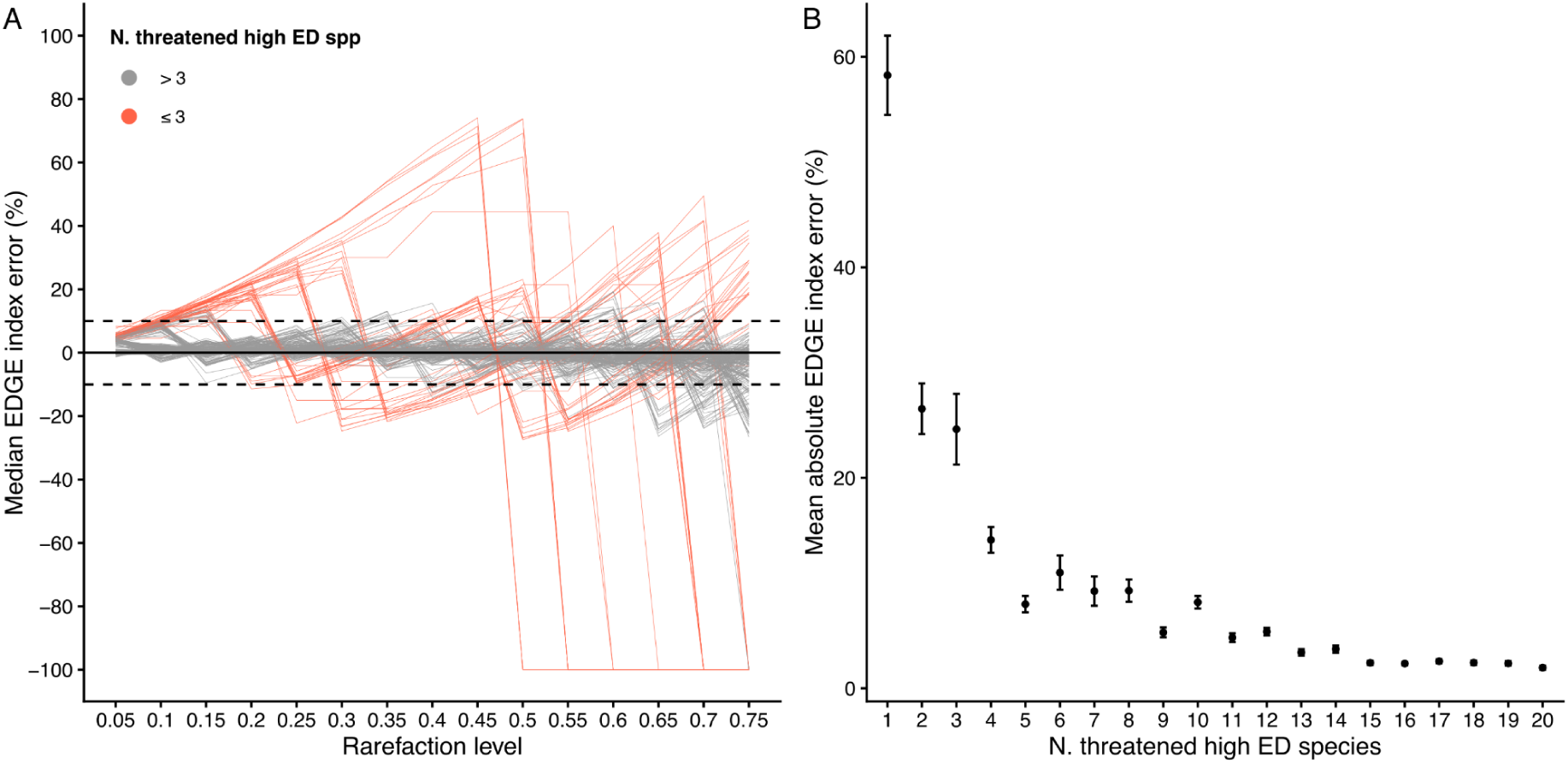
The EDGE Index error under rarefaction for birds. A: Median EDGE Index error across rarefaction levels, stratified by whether a country has more (grey) or less than or equal to (red) three threatened high ED bird species. Lines reflect individual countries, with dashed horizontal lines indicating ±10% error, plotted as a visual reference. B: Mean absolute EDGE Index error across all rarefaction levels in countries grouped by their number of high ED threatened bird species, with uncertainty bars indicating the standard error. Abbreviation: N. threatened high ED spp = Number of threatened high ED species.

From this finding, we can make a general recommendation for those seeking to report a country-level EDGE Index. Namely, in countries where there are three or fewer high ED threatened species present for a given taxonomic group, EDGE Index can be made more robust by aggregating results from across taxonomic groups. For example, rather than reporting on tortoises & turtles, lizards & snakes, and crocodylians individually, when the number of high ED threatened species in each is low, it would be more robust to report on reptiles as a whole.

## 7. Applications & Outlook - Perspectives

Recent global analyses on the reporting of KM-GBF indicators have found that few countries are as of yet reporting on component and complementary indicators (https://s3.amazonaws.com/cbddocumentspublic-imagebucket-15w2zyxk3prl8/a61b896636c945199136ea9bef08b78b-3). Although not compulsory like headline indicators, component and complementary indicators are still important because, for most KM-GBF targets, there is no single indicator that can measure whether or not a target has been achieved. For instance, the RLI is the headline indicator for Target 4 but in overlooking the phylogenetic dimension of the biodiversity crisis, the biodiversity targets fail to represent the elevated extinction risks of ED species within some groups, which the EDGE Index fully accounts for. Furthermore, many of these component and complementary indicators also lack methodological guidance (https://www.gbf-indicators.org/complementary-indicators), which may act as a barrier to their uptake. In this study, we have established clear guidance on how to calculate the country-level EDGE Index, using either national or global Red List data, which can facilitate the monitoring of high ED species at a national level and promote the reporting of the country-level EDGE Index more broadly.

### 7.1. On the importance of using national *vs.* global Red List data for the country-level EDGE Index

Our results show that the National EDGE Index differs from the Global EDGE Index in both French and South African practical applications, with nationally derived indices usually showing elevated national threat levels. By using national extinction risk data, the National EDGE Index better reflects local priorities, given that nationally produced Red List assessments reflect specific threats to biodiversity within country borders, rather than global threats. This approach should be more valuable to countries assessing their existing measures to safeguard their ED species, and for informing decisions on which species to prioritise for conservation actions. This is consistent with general trends whereby other types of national indicators, such as the RLI and forest cover loss, differ from globally derived versions, although these differences are not always consistent in their magnitude or direction (Han et al., 2017).

There also remains a strong interest in applying global indicators at a national level produced following standardised procedures. Indeed, they can reveal general trends for the same sets of species (Butchart et al., 2025), and thus facilitate the global monitoring of biodiversity, whilst ensuring comparability between countries (Buchanan et al., 2020). For the EDGE Index, globally derived values have been used here to reveal stark spatial differences in the proportions of threatened high ED species for a variety of taxonomic groups of jawed vertebrates. By comparison, only half of nations have national extinction risk data, with varying degrees of comprehensiveness and varied taxonomic focuses (Raimondo et al., 2023). Furthermore, National Red Lists can follow different criteria from the IUCN (Regan et al., 2005), and some, like the United States Endangered Species Act, can display marked differences from the IUCN Red List (Harris et al., 2012). Despite these discrepancies, the global threat status of non-endemic species will typically fail to reflect their national threat status, and building and maintaining taxonomically broad and comprehensive National Red Lists across all countries remains immensely valuable for the global conservation of biodiversity. Then, the National EDGE Index accounts for local threats that may not be apparent in the global assessments, as well as local conservation efforts that may ameliorate extinction risk nationally despite high threats elsewhere in the range of a species. There is thus a utility in calculating and reporting both national and global values of a biodiversity indicator and any difference between the two is informative; for the EDGE Index, a difference can frame local extirpation risks of high ED species in a finer context of their global extinction risks, and could serve to galvanise conservation efforts at any sub-global level.

### 7.2. Perspectives

The taxonomic scope of the EDGE Index is currently limited to the taxonomic groups for which phylogenetic and extinction risk data are available to estimate evolutionary distinctiveness of species at a global scale. Parties of CBD agreed that threat assessments should become more frequent, at least for the major taxonomic groups. At the same time, big phylogenetic trees are becoming available for several very species-rich groups such as plants (*e.g.*, flowering plants, Carruthers et al., 2026) and invertebrates (*e.g.*, dragonflies and damselflies, Bybee et al., 2021). Although some major taxonomic groups such as fungi (Cunha et al., 2026) lack the requisite data needed for such assessments, alternative approaches are being developed to cover a larger proportion of the tree of life. For example, when it comes to determining the evolutionary distinctiveness of species, predictive methods are facilitating a greater utilisation of Open Tree of Life data (Duke et al., 2026), which has been used to generate predicted evolutionary distinctiveness for roughly 2.2 million species (Guo et al., 2025). Meanwhile, taxonomic approaches, which can highlight species from monotypic genera or families (Gumbs et al., 2024), are a useful proxy for identifying candidate high ED species in data poor clades like fungi (Cunha et al., 2026). Predictive modelling of extinction risk has been employed successfully for hyper-diverse taxonomic groups like flowering plants (Bachman et al., 2024) and ray-finned fish (Chen et al., 2023); predictive methods have been utilised in recent EDGE calculations (Forest et al., 2026).

An increasing number of global conservation assessments are also establishing trend baselines for diverse groups, such as for dragonflies and damselflies (Sayer et al., 2025), sharks & rays (Dulvy et al., 2021), and reptiles (Cox et al., 2022), and sampled approaches provide an effective alternative for taxonomic groups that would otherwise be neglected (Butchart et al., 2025). The IUCN Red List framework leads to the deployment of incredible resources to generate data that lies at the heart of the EDGE approach and in particular for the EDGE Index. This growing wealth of data should therefore enable EDGE Index calculations for a much broader array of taxonomic groups in the future. Furthermore, it is all the more important to strengthen efforts to accumulate knowledge to reduce the number of species assessed as DD/NE, as this has an impact on the value and reliability of the EDGE Index of a given taxonomic group in a given spatial unit (see sections 3., 4.; Figures 2 and 3). More broadly, threat assessments can facilitate the protection of evolutionary history in national policies and local conservation efforts. The more data we have available, the more we can define appropriate, effective and relevant protection actions for the conservation of biodiversity including preserving phylogenetic diversity, one of its essential facets not sufficiently captured through species-centred measures (Burgess et al., 2024).

## 8. Conclusion

Parties of the CBD have committed to building and maintaining National Red Lists, which represent the bare minimum in terms of tracking national biodiversity conservation efforts. Additionally, a growing number of global phylogenetic trees are available thanks to advances in sequencing and phylogenetic methods. Therefore, calculating evolutionary distinctiveness at a global scale is becoming possible for many groups across the tree of life, with the PDSG making this data centralised and openly available. Such advances, including the method we propose here, facilitate the calculation of the EDGE Index at a national level and for many countries.

As an indicator monitoring the changing conditions of the tree of life featured in the KM-GBF, the EDGE Index enshrines the importance of conserving evolutionary history within an internationally agreed framework. The protocol outlined here for incorporating national extinction risk assessments into the country-level EDGE Index will help to ensure that evolutionarily distinct species across the world are monitored and conserved, and all the more effectively when conservation efforts are tailored to local threat levels.

## Contributions

Project conceptualisation and design: all co-authors.

Generated datasets: JLMC, CR for France (section 3.); KAT, JANL for South Africa (section 4.); SP, RG for global data (section 5.).

Script conception: SP, JLMC for the general methods of calculation and compilation; JLMC for France (section 3.); JANL for South Africa (section 4.); SP for global disaggregation (section 5.) and robustness (section 6.).

Formal analysis: JLMC for France (section 3.); JANL, KAT for South Africa (section 4.); SP for global data (section 5.) and robustness (section 6.).

Investigation: JLMC, RP for France (section 3.); KAT, JANL for South Africa (section 4.); SP, JLMC, RG for global data (section 5.) and robustness (section 6.).

Writing—original draft: JLMC, SP for Introduction (section 1.), Methods (section 2.), Discussion & Perspectives (section 7.), Conclusion (section 8.); JLMC for France (section 3.); KAT, JANL for South Africa (section 4.); SP for global data (section 5.) and robustness (section 6.).

Writing—review and editing: JLMC, SP, JANL, FF, RP, KAT, RG. Final review: all co-authors.

## Type of article

Contributed paper

## Single sentence that describes the target audience for the paper

The article targets the researchers, conservation practitioners, and policy advisors who wish to monitor the effectiveness of national conservation efforts to safeguard the tree of life.

## Acknowledgements

We thank Audrey Delhomme for facilitating the organization of the workshop “Biodiversity On the EDGE”. We also thank Dr. Carol Poole for her support on this work.

## Data Accessibility Statement

### Ethics approval statement

Not applicable

