## Supplementary materials for "A framework for measuring national trends in the extinction risk of evolutionarily distinct species using the EDGE Index"

**Title**

( ◇ equal first authors | ♦ equal lead authors)

**Affiliations**

[1]*Institut de Systématique, Evolution, Biodiversité (ISYEB), Muséum national d'Histoire naturelle, CNRS, SU, EPHE-PSL, UA, 57 rue Cuvier, CP50, 45 rue Buffon, 75005 Paris, France*

[2]*PatriNat, Muséum national d'Histoire naturelle, Office Français de la Biodiversité, CNRS, IRD, Maison Buffon, CP41, 36 rue Geoffroy Saint-Hilaire, 75005 Paris.*

[3]*Royal Botanic Gardens, Kew, Richmond, UK*

[4]*Zoological Society of London, London, UK*

[5]*Imperial College London, UK*

[6]*Studio BlueGreen, London, UK*

[7]*Science and Solutions for a Changing Planet Doctoral Training Programme, Grantham Institute, London, UK*

[8]*Universidad Autónoma de Madrid, Spain*

[9]*South African National Biodiversity Institute, South Africa*

[10]*Department of Zoology, Centre for Ecological Genomics and Wildlife Conservation, University of Johannesburg, P.O. Box 524, Johannesburg, South Africa*

[11]*Anura Africa, Durban, South Africa*

[12]*Institute for Coastal and Marine Research, PO Box 77000, Nelson Mandela University, Gqeberha, 6031, South Africa*

### Practical application: French EDGE Index for nesting birds across ten territories

#### Material

##### 1) Description of the datasets

TAXREF is the national taxonomic and nomenclatural register of reference for species living in Metropolitan and Overseas France, built and updated by Museum National d'Histoire Naturelle (MNHN) in the application framework of the Système d'Information de l'iNventaire du Patrimoine naturel (SINP). It aims to centralize and organize information on the species present in the different French territories notably with their most recent taxonomy, synonyms included, and their ecological environment such as continental *vs.* marine. For a given species, its validated scientific name and associated synonyms are considered at the light of the taxonomic knowledges at a given time and according to the specific nomenclatural codes as following:

- ICZN for animals (https://www.iczn.org/the-code/the-code-online/) ;
- Shenzhen code for algaes, fungi and plants (https://www.iapt-taxon.org/nomen/main.php) ;
- procaryote code for bacteria (https://www.microbiologyresearch.org/content/journal/ijsem/10.1099/ijsem.0.000778).

Different sources are also used to complete and/or update TAXREF such as several Global Species Database of reference (*e.g.*, WoRMS, Index Fungorum), scientific publications or data from expertise groups (*e.g.*, Société Herpétologique de France) but also Regional Species Database, Local Species Database and Global Biodiversity Information Facility (GBIF; https://www.gbif.org/). Inversely, TAXREF contributes to providing national data for Species Databases or GBIF.

TAXREF focuses primarily on cataloging the wild species that have been present in France since the Paleolithic era. Domestic or cultivated taxa are not included, except under some conditions such as the monitoring of invasive species. The taxonomic status of species such as "species" in the biological sense, "subspecies", or even "variety" are also differentiated.

TAXREF also aims at inventorying the geographical distribution of species across French territories through the use of biogeographic status per administrative spatial unit. The range of biogeographic status indicated considers three main concepts: (1) presence or absence and even extinction status, (2) origin (native or alien/introduced), and (3) occupied area (endemism). The errors of species observation are also considered facilitating the updates between the different versions of TAXREF. The attribution of biogeographic status is based on scientific publications. More details are provided in Gargominy et al. 2022. For this study, we used the data from version 18 of TAXREF published in 2025 (Gargominy et al. 2025) – hereafter TAXREFv18– and provided by INPN from https://inpn.mnhn.fr/referentiels-donnees. The key features of French biodiversity across taxonomic groups are provided below in Table S1.

BDC-Statuts (Base De Connaissance) is the dataset cataloging the information relative to the status of species present in France in the different programs such as global and national IUCN assessments, regulations on species protection at global, european, national, territorial scales, etc. More details are provided in Régnier & Gargominy  2018. For this study, we used the data from version 18 of BDC-Statuts published in 2025 –hereafter BDC-Statutsv18– and provided by INPN from https://inpn.mnhn.fr/referentiels-donnees. The key features of the number of threatened species across taxonomic groups are provided below in Table SX1

**Table S1. Key features regarding French biodiversity and number of threatened species across taxonomic groups**. Calculations of the different key features are adapted from the methodology used in Barnier et al. 2026. NB: Metropolitan France includes Corsica island located in the Mediterranean Sea.

| Territory | Configuration | Location | Number of species | Number of threatened species |
|---|---|---|---|---|
| Metropolitan France | Continental | Europe - Mediterranean Sea | 108,825 | 1,181 |
| French Guiana | Continental | South America - Atlantic Ocean | 31,258 | 298 |
| Martinique | Insular | Caribbean Sea | 8,867 | 368 |
| Guadeloupe | Insular | Caribbean Sea | 12,540 | 454 |
| Saint-Martin | Insular | Caribbean Sea | 2,477 | 117 |
| Saint-Barthélémy | Insular | Caribbean Sea | 2,396 | 115 |
| Saint-Pierre-et-Miquelon | Insular | Atlantic Ocean | 2,927 | 89 |
| Mayotte | Insular | Indian Ocean | 6,970 | 405 |
| Scattered islands | Insular | Indian Ocean | 2,265 | 114 |
| La Réunion | Insular | Indian Ocean | 13,688 | 568 |
| Subantarctic islands | Insular | Indian & Austral Oceans | 3,527 | 50 |
| Adélie Land | Continental | Antarctic - Austral Ocean | 1,033 | 9 |
| French Polynesia | Insular | Pacific Ocean | 13,861 | 463 |
| New Caledonia | Insular | Pacific Ocean | 31,438 | 834 |
| Wallis-et-Futuna | Insular | Pacific Ocean | 3,052 | 99 |
| Clipperton | Insular | Pacific Ocean | 977 | 22 |

To build the the national ED species list for France, we first made a taxonomic reconciliation between the global ED data from the PDSG platform, the taxonomic backbone from IUCN Red List of 2025 to check the updates of binomial names for a given Red List identifier, and with TAXREFv18 (STEP2, blue patch in Figure 2). Then, we merge the French ED species list with BDCv18 through both Red List identifier and the TAXREFv18 identifier for a given scientific name to obtain the final dataset used for the practical application for France (STEP3 →4, blue patch in Figure 2).

Hence, the final dataset used for the practical application of French EDGE Index was composed of species with the following characteristics:

- from TAXREFv18, with the taxonomic status: species (ES), subspecies (SSES) for which subspecies were the only representative of a species on a given territory and considered equivalent to the related species (see above Table S2) ;
- from TAXREFv18, with the biogeographical status on the specific territory as: present (P), endemic (E), subendemic (S), native (N), locally extinct in the wild but not globally (W), globally extinct (X) and endemic extinct (Z);
- from BDC-Statutsv18 (and IUCN France publications for the studied territories), with the seasonal status of nesting species;
- from the taxonomic reconciliation with the global ED dataset, species matching was comprehensive (see below).

The final dataset used for the two versions of the French EDGE Index compilations and the different analyses is provided here, and the summary of the French EDGE Index results for the ten territories calculated using National Red List and IUCN Red List is provided below in Table S3.

#### 2) Presentation of the characteristics of the ten French territories used in this study

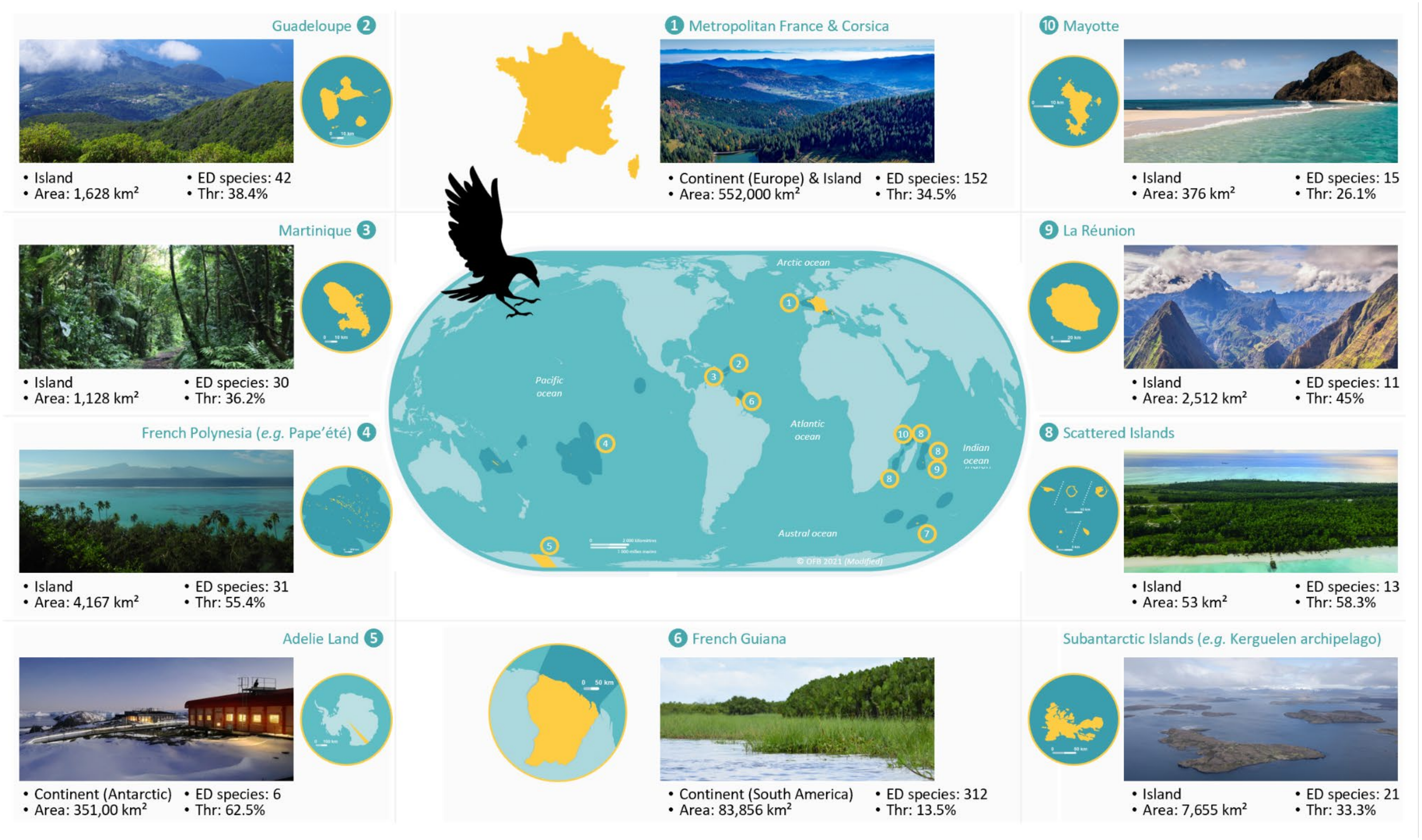

**Figure S1**. **Presentation of the main characteristics of the ten French territories used in this study** with their geographical location, the kind of territory (island *vs.* continental), their area (km²), the number of high ED species of nesting birds present in the territory (ED species), the percentage of species assessed as threatened in the territory (Thr). Photo credits: Guadeloupe, Parc National ©O.Delzons; Martinique, Le Morne Rouge ©Guillaume Gigot; French Polynesia, Tahiti © Guillaume Gigot; Adelie Land, Durmont d'Urville Station ©TAAF; French Guiana, De Raw-Roura marsh ©J.-P. Siblet; Subantarctic islands, Kerguelen archipelago ©Bruno Maire; Scattered islands ©Stéphanie Legeron; La Réunion, Cap Noir ©Adobestock/A.Karnholz; Mayotte, Choizil islet ©Adobestock/Michel Bordieu; Metropolitan France including Corsica, trout lake in the Vosges mountains ©Adobestock/Dominique.

**Table S2. List of the subspecies that were the only representative of a species on a given territory and considered equivalent to the related species.**

| Territory | Subspecies | Species |
|---|---|---|
| Martinique | *Setophaga petechia ruficapilla* | *Setophaga petechia* |
| French Guiana | *Myiodynastes maculatus maculatus* | *Myiodynastes maculatus* |
| French Guiana | *Progne chalybea chalybea* | *Progne chalybea* |
| French Guiana | *Progne tapera tapera* | *Progne tapera* |
| French Guiana | *Thalasseus sandvicensis eurygnathus* | *Thalasseus sandvicensis* |
| Mayotte | *Accipiter francesii brutus* | *Accipiter francesii* |
| Mayotte | *Alectroenas sganzini sganzini* | *Alectroenas sganzini* |
| Mayotte | *Apus barbatus mayottensis* | *Apus barbatus* |
| Mayotte | *Bubulcus ibis ibis* | *Bubulcus ibis* |
| Mayotte | *Butorides striata rhizophorae* | *Butorides striata* |
| Mayotte | *Corythornis vintsioides johannae* | *Corythornis vintsioides* |
| Mayotte | *Cypsiurus parvus griveaudi* | *Cypsiurus parvus* |
| Mayotte | *Foudia eminentissima algondae* | *Foudia eminentissima* |
| Mayotte | *Gallinula chloropus pyrrhorrhoa* | *Gallinula chloropus* |
| Mayotte | *Leptosomus discolor discolor* | *Leptosomus discolor* |
| Mayotte | *Nesoenas picturata comorensis* | *Nesoenas picturata* |
| Mayotte | *Tachybaptus ruficollis capensis* | *Tachybaptus ruficollis* |
| Mayotte | *Terpsiphone mutata pretiosa* | *Terpsiphone mutata* |
| Mayotte | *Tyto alba affinis* | *Tyto alba* |
| Mayotte | *Zosterops maderaspatanus mayottensis* | *Zosterops maderaspatanus* |
| Scattered islands | *Phaethon lepturus europae* | *Phaethon lepturus* |
| Scattered islands | *Puffinus bailloni bailloni* | *Puffinus bailloni* |
| Scattered islands | *Sula dactylatra melanops* | *Sula dactylatra* |
| La Réunion | *Gallinula chloropus pyrrhorrhoa* | *Gallinula chloropus* |
| La Réunion | *Psittacula eques eques* | *Psittacula eques* |
| La Réunion | *Puffinus lherminieri bailloni* | *Puffinus bailloni* |
| La Réunion | *Terpsiphone bourbonnensis bourbonnensis* | *Terpsiphone bourbonnensis* |
| La Réunion | *Zosterops borbonicus borbonicus* | *Zosterops borbonicus* |
| Subantarctic islands | *Larus dominicanus judithae* | *Larus dominicanus* |

#### Results

**Table S3. Summary of the French EDGE Index results for the ten territories calculated using National Red List and IUCN Red List.** Configuration: indicates if the territory presents a continental *vs.* insular topography; Year: date of the national Red List assessment; N.ass sp: number of assessed species. N.sp DD/NE: number of species categorized as DD/NE; N.ED sp (% tot sp): number of high ED species (percentage relatively to the total number of species, *i.e.*, N.ass sp + N.sp DD/NE); N.ED sp and ass (% N.ass. sp): number of high ED species and assessed (percentage relatively to the number of assessed species, *i.e.*, N.ass sp); N.ED sp DD/NE (%N.sp DD/NE): number of high ED species categorized DD/NE (percentage relatively to the number of species categorized as DD/NE, *i.e.*, N.sp DD/NE) with meanNRL = 8.81% *vs.* meanGRL = 1.16%; EDGE Index [confidence interval]: value of EDGE Index with its associated confidence interval calculated by using the number of DD/NE species relative to the National Red List.

| Territory | Configuration | Year | N.ass sp | | N.sp DD/NE | | N.ED sp (% tot sp) | |
|---|---|---|---|---|---|---|---|---|
| | | | National | Global | National | Global | National | Global |
| Metropolitan France | Continental | 2016 | 276 | 270 | 2 | 8 | 152 (54.7%) | 152 (54.7%) |
| Guadeloupe | Insular | 2021 | 66 | 73 | 7 | 0 | 42 (57.5%) | 42 (57.5%) |
| Martinique | Insular | 2020 | 57 | 57 | 1 | 1 | 30 (51.7%) | 30 (51.7%) |
| French Guiana | Continental | 2017 | 494 | 553 | 61 | 2 | 312 (56.2%) | 312 (56.2%) |
| French Polynesia | Insular | 2015 | 56 | 56 | 0 | 0 | 31 (55.4%) | 31 (55.4%) |
| Mayotte | Insular | 2014 | 22 | 22 | 1 | 1 | 15 (65.2%) | 15 (65.2%) |
| Scattered Islands | Insular | 2015 | 12 | 20 | 8 | 0 | 13 (65%) | 13 (65%) |
| La Réunion | Insular | 2010 | 20 | 19 | 0 | 1 | 11 (55%) | 11 (55%) |
| Subantarctic Islands | Insular | 2015 | 36 | 43 | 9 | 2 | 21 (46.7%) | 21 (46.7%) |
| Adélie Land | Continental | 2015 | 7 | 8 | 1 | 0 | 6 (75%) | 6 (75%) |

**Table S3.** continued

| Territory | Configuration | Year | N.ED sp and ass (% N.ass sp) | | N.ED sp DD/NE (% N.sp DD/NE) | | EDGE Index [confidence intervall] | |
|---|---|---|---|---|---|---|---|---|
| | | | National | Global | National | Global | National | Global |
| Metropolitan France | Continental | 2016 | 151 (99.3%) | 150 (98.7%) | 1 (0.66%) | 2 (1.32%) | 0.344 [0.342, 0.349] | 0.013 [0.013, 0.020] |
| Guadeloupe | Insular | 2021 | 37 (88.1%) | 42 (100%) | 5 (11.9%) | 0 (0%) | 0.486 [0.429, 0.548] | 0.048 [0.048, 0.048] |
| Martinique | Insular | 2020 | 29 (96.7%) | 29 (96.7%) | 1 (3.33%) | 1 (3.33%) | 0.379 [0.367, 0.400] | 0.034 [0.033, 0.067] |
| French Guiana | Continental | 2017 | 276 (88.5%) | 311 (99.7%) | 36 (11.5%) | 1 (0.32%) | 0.178 [0.163, 0.192] | 0.003 [0.003, 0.006] |
| French Polynesia | Insular | 2015 | 31 (100%) | 31 (100%) | 0 (0%) | 0 (0%) | 0.581 [0.581, 0.581] | 0.452 [0.452, 0.452] |
| Mayotte | Insular | 2015 | 14 (93.3%) | 14 (93.3%) | 1 (6.67%) | 1 (6.67%) | 0.357 [0.333, 0.400] | 0.143 [0.133, 0.200] |
| Scattered Islands | Insular | 2015 | 10 (76.9%) | 13 (100%) | 3 (23.1%) | 0 (0%) | 0.600 [0.462, 0.692] | 0.154 [0.154, 0.154] |
| La Réunion | Insular | 2010 | 11 (100%) | 11 (100%) | 0 (0%) | 0 (0%) | 0.455 [0.455, 0.455] | 0.182 [0.182, 0.182] |
| Subantarctic Islands | Insular | 2015 | 18 (85.7%) | 21 100%) | 3 (14.3%) | 0 (0%) | 0.389 [0.333, 0.476] | 0.143 [0.143, 0.143] |
| Adélie Land | Continental | 2015 | 5 (83.3%) | 6 (100%) | 1 (16.7%) | 0 (0%) | 0.600 [0.500, 0.667] | 0 [0, 0] |

### Practical application: South African EDGE Index for jawed vertebrates with a particular attention to endemic amphibians, lizards & snakes, tortoises & turtles

#### Results

The EDGE Index was calculated temporally for South Africa's herpetofauna. Red list categories were used from 1990, 2016, and 2022 for lizards & snakes and tortoises & turtles, and 2004, 2016, and 2025 for amphibians. Non-genuine changes—those resulting from factors such as updated geographic information or taxonomic revisions that would have affected earlier assessments—were accounted for by applying the species' current threat category to previous assessment years (*i.e.*, backcasting). Subsequently, lists of EDGE species were generated for amphibians and reptiles (Table S4).

**Table S4. Lists of EDGE a) amphibians and b) reptiles and their threat status (RL) over multiple assessment time points.** Indicated are the scores for evolutionary distinctiveness (ED), derived from a global phylogeny (Gumbs et al., 2024), and the weighted score incorporating threat status as an extinction probability (EDGE). All South African amphibians are in the Order Anura. Species marked with an asterisk (*) did not appear in the global phylogeny; therefore, an average ED score for the genus was used. Marine testudines were not included.

| **a)** | | | **Red List Status** | | | | **Scores** | |
|---|---|---|---|---|---|---|---|---|
| **Family** | **Species** | **South African Category** | **2004** | **2016** | **2025** | **Global** | **ED** | **EDGE** |
| *Pyxicephalidae* | *Cacosternum platys* | Endemic | EN | EN | EN | NT | 34.86 | 24.60 |
| Heleophrynidae | *Heleophryne rosei* | Endemic | EN | EN | EN | CR | 19.87 | 13.92 |
| Heleophrynidae | *Heleophryne hewitti* | Endemic | EN | EN | EN | EN | 18.95 | 13.18 |
| Pyxicephalidae | *Microbatrachella capensis* | Endemic | EN | EN | EN | CR | 16.36 | 11.48 |
| *Arthroleptidae* | *Arthroleptis wageri* | Endemic | EN | EN | EN | NA | 15.63 | 10.93 |
| Pyxicephalidae | *Arthroleptella rugosa* | Endemic | CR | CR | CR | CR | 11.82 | 10.67 |
| Pipidae | *Xenopus gilli* | Endemic | EN | EN | EN | EN | 14.62 | 10.28 |
| Pyxicephalidae | ***Cacosternum cederbergense** * | Endemic | EN | EN | EN | NA | 13.37 | 9.32 |
| Pyxicephalidae | *Anhydrophryne rattrayi* | Endemic | EN | EN | EN | VU | 12.93 | 9.00 |
| Brevicipitidae | *Breviceps bagginsi* | Endemic | CR | CR | CR | NT | 9.16 | 8.26 |
| Pyxicephalidae | *Anhydrophryne ngongoniensis* | Endemic | EN | EN | EN | EN | 11.79 | 8.23 |
| Pyxicephalidae | *Cacosternum thorini* | Endemic | EN | EN | EN | EN | 9.67 | 6.73 |
| Hyperoliidae | *Afrixalus knysnae* | Endemic | EN | EN | EN | EN | 9.27 | 6.50 |
| Brevicipitidae | ***Breviceps batrachophiliorum** * | Endemic | EN | EN | EN | NA | 9.23 | 6.47 |
| Arthroleptidae | *Leptopelis xenodactylus* | Endemic | EN | EN | EN | EN | 8.51 | 5.98 |
| Pyxicephalidae | *Amietia hymenopus* | Not Endemic | NT | NT | VU | NT | 11.87 | 5.93 |
| Pyxicephalidae | *Cacosternum capense* | Endemic | VU | VU | VU | NT | 11.12 | 5.59 |
| Pyxicephalidae | *Cacosternum aggestum* | Endemic | VU | VU | VU | LC | 11.03 | 5.55 |
| Pyxicephalidae | *Arthroleptella bicolor* | Endemic | VU | VU | VU | NT | 10.43 | 5.19 |
| Brevicipitidae | *Breviceps sylvestris* | Endemic | VU | VU | VU | NT | 9.07 | 4.53 |
| Brevicipitidae | *Breviceps gibbosus* | Endemic | VU | VU | VU | NT | 8.43 | 4.16 |

CR-Critically Endangered, EN-Endangered, VU-Vulnerable, NT-Near Threatened, LC-Least Concern, NA - Not Assessed

| **b)** | | | | **Red List Status** | | | | **Scores** | |
|---|---|---|---|---|---|---|---|---|---|
| **Order** | **Family** | **Species** | **South African Category** | **1990** | **2018** | **2022** | **Global** | **ED** | **EDGE** |
| Squamata | Cordylidae | *Smaug giganteus* | Endemic | VU | VU | VU | VU | 38 | 18.8875 |
| Testudines | Testudinidae | *Psammobates geometricus* | Endemic | EN | CR | CR | CR | 13 | 11.6925 |
| Testudines | Testudinidae | *Chersobius signatus* | Endemic | NT | EN | EN | EN | 13 | 9.0557 |
| Squamata | Cordylidae | *Hemicordylus nebulosus* | Endemic | VU | VU | VU | VU | 15 | 7.4399 |
| Squamata | Scincidae | *Scelotes inornatus* | Endemic | EN | CR | CR | CR | 7 | 6.2974 |
| Squamata | Scincidae | *Scelotes kasneri* | Endemic | NT | NT | EN | EN | 8 | 5.5874 |
| Squamata | Chamaeleonidae | *Bradypodion caeruleogula* | Endemic | EN | EN | EN | EN | 8 | 5.5867 |
| Squamata | Scincidae | *Scelotes bourquini* * | Endemic | VU | VU | VU | NA | 8.35 | 4.2161 |

CR-Critically Endangered, EN-Endangered, VU-Vulnerable, NT-Near Threatened, LC-Least Concern, NA - Not Assessed

### Practical application: Global EDGE Index for all jawed vertebrates

#### Results

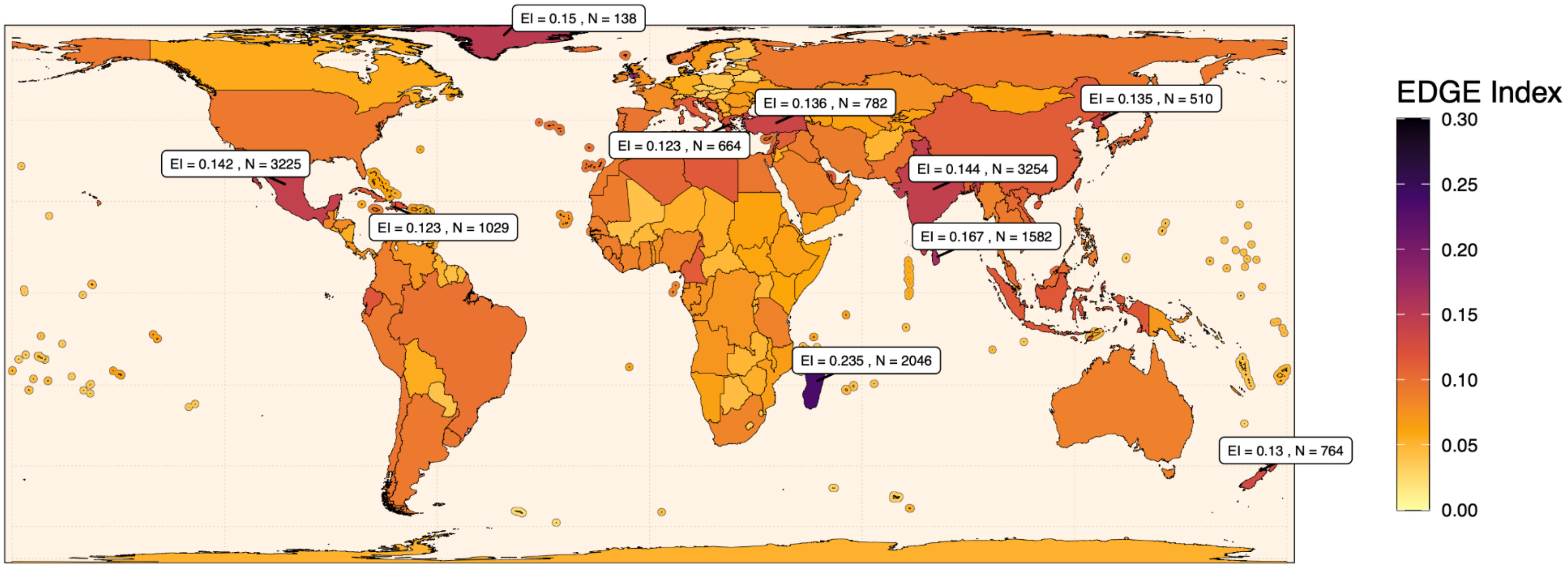


**Figure S2. Global EDGE Index aggregated for all jawed vertebrates using IUCN Red List data of 2023.** Labels indicate the EDGE Index (EI) and the number of high ED species (N) in the ten highest ranked countries for their EDGE Index.

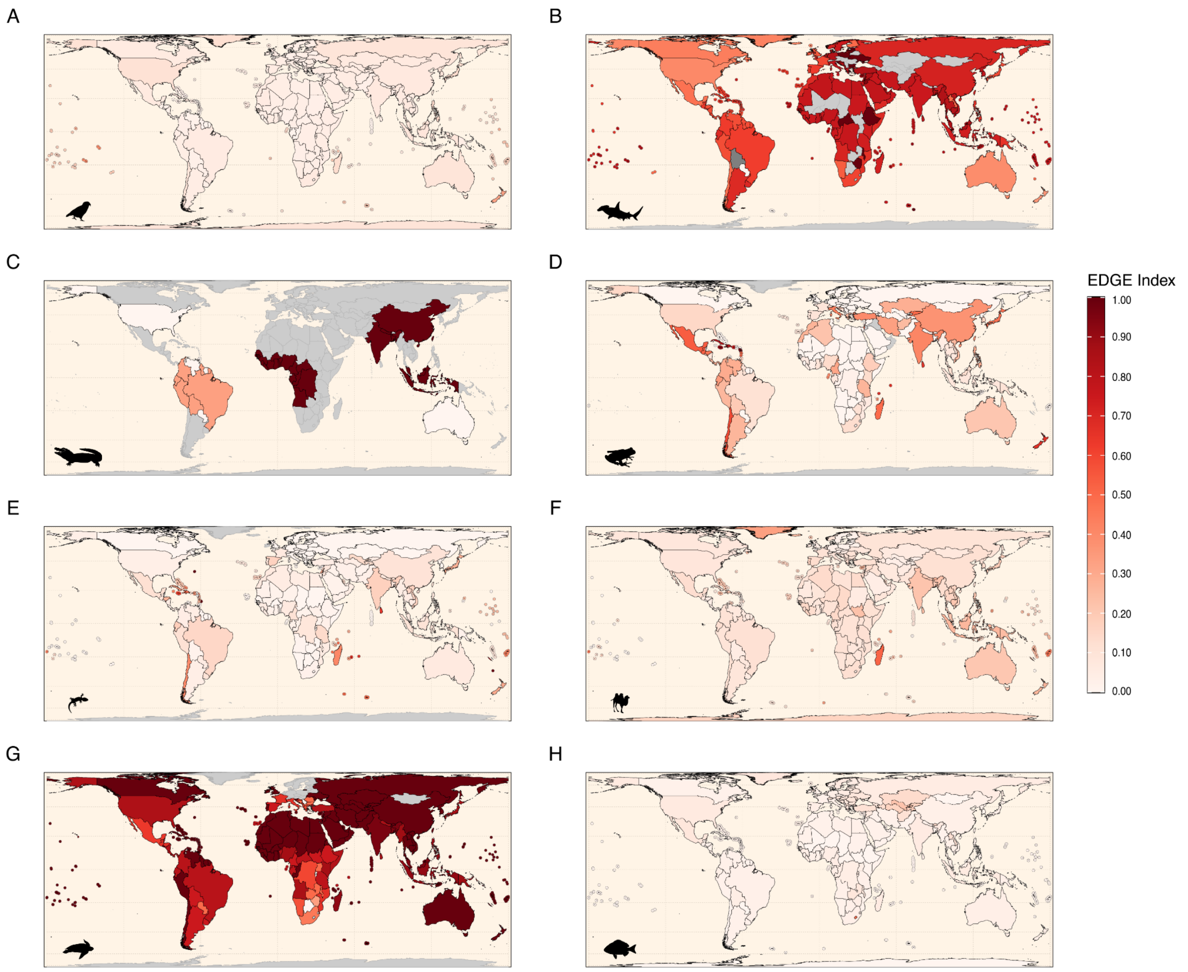


**Figure S3. Global EDGE Index for all taxonomic groups of jawed vertebrates (individually) using IUCN Red List data of 2023.** From A-H: birds, sharks & rays, crocodylians, amphibians, lizards & snakes, mammals, tortoises & turtles, and ray-finned fish. Grey spatial units indicate regions with no high ED species.

### EDGE Index robustness

#### Results

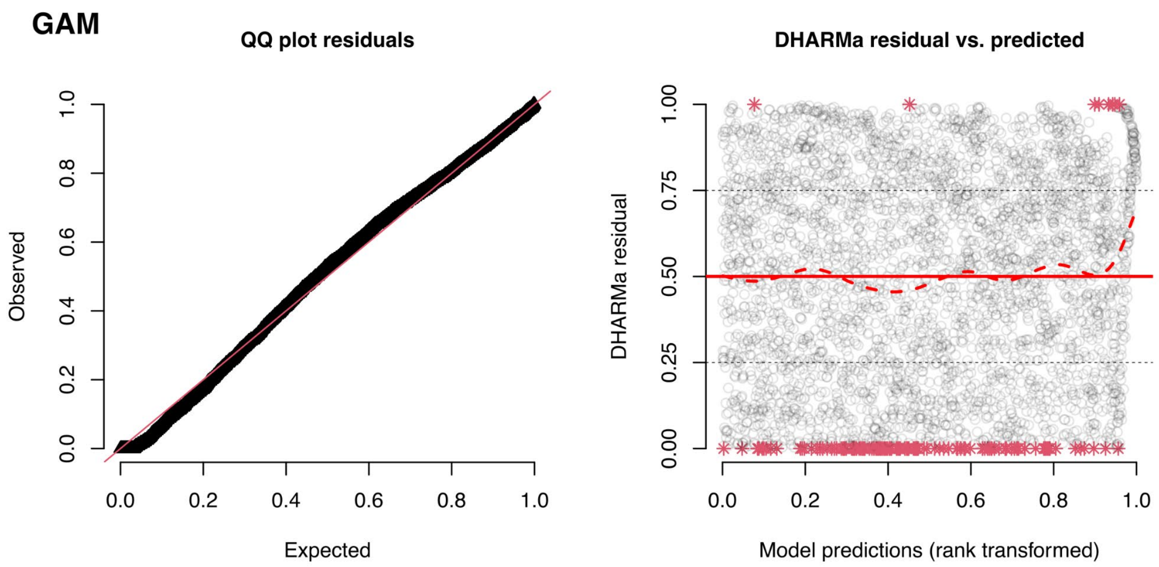


**Figure S4. Model diagnostics of generalised linear mixed model fit to Global EDGE Index error.** The GAMM used a log-link function and a gamma distribution and included the following fixed effects: the number of threatened high ED species, the number of high ED species, and the rarefaction level. Country was included as a random intercept. Richness measures were modelled as smooth terms. The left panel shows QQ plots, used to assess the normality of scaled residuals. The right panel shows scaled residuals *vs.* predicted values, used to assess homoscedasticity. Diagnostic plots were made using the DHARMa package in R (Hartig and Hartig, 2017), which produces simulated residuals based on the fitted model.

**Table S5. Coefficient table for a generalised additive mixed model evaluating the effects of richness and rarefaction on Global EDGE Index error in birds.** Rarefaction level was modelled as a parametric fixed effect. s(N.tED) and s(N.ED) indicate the number of threatened high ED bird species and the number of non-threatened high ED bird species, respectively, both modelled as smooth terms. s(country) refers to the country, modelled as a random intercept. The column e.d.f refers to the estimated degrees of freedom.

| | **Estimate** | **Std. Error** | **t-value** | **p-value** |
|---|---|---|---|---|
| **(Intercept)** | 0.618 | 0.032 | 19.612 | < 0.001 |
| **Rarefaction level** | 1.209 | 0.059 | 20.573 | < 0.001 |

| | **e.d.f.** | | **F** | **p-value** |
|---|---|---|---|---|
| **s(N.tED)** | 6.784 | | 220.955 | < 0.001 |
| **s(N.ED)** | 5.829 | | 12.421 | < 0.001 |
| **s(country)** | 144.896 | | 3.097 | < 0.001 |
